# Monocyte recruitment to the inflamed central nervous system: migration pathways and distinct functional polarization

**DOI:** 10.1101/2020.04.04.025395

**Authors:** Daniela C. Ivan, Sabrina Walthert, Giuseppe Locatelli

## Abstract

The central nervous system (CNS) parenchyma is enclosed by anatomical interfaces including multilayered meninges, the blood-brain barrier (BBB), the choroid plexuses within ventricles and the glia limitans. These border areas hold distinct functional specializations which control the trafficking of monocyte-derived cells toward the CNS parenchyma, altogether maintaining CNS homeostasis. By crossing activated endothelial, epithelial and glial borders, circulating leukocytes gain however access to the CNS parenchyma in several inflammatory diseases including multiple sclerosis.

Studies in animal models of neuroinflammation have helped describing the phenotypic specifications of these invading monocyte-derived cells, able to exert detrimental or beneficial functions depending on the local environment. In this context, *in vivo* visualization of iNOS^+^ pro-inflammatory and arginase-1^+^ anti-inflammatory macrophages has recently revealed that these distinct cell phenotypes are highly compartmentalized by CNS borders. While arginase-1^+^ macrophages densely populate the leptomeninges, iNOS^+^ macrophages rather accumulate in perivascular spaces and at the pia mater-CNS parenchymal interface.

How and where these macrophages acquire their functional commitment, and whether differentially-activated monocyte-derived cells infiltrate the CNS through distinct gateways, remains however unclear.

In this study, we have investigated the interaction of monocyte-derived macrophages with endothelial (BBB) and epithelial (choroid plexus) barriers of the CNS, both *in vitro* and *in vivo*. By using primary mouse brain microvascular endothelial cells as *in vitro* model of the BBB, we observed that, compared to unpolarized primary macrophages, adhesion of functionally-committed macrophages to endothelial cells was drastically reduced, literally abrogating their diapedesis across the BBB. Conversely, when interacting with an activated choroid plexus epithelium, both pro- and anti-inflammatory macrophages displayed substantial adhesive and migratory properties. Accordingly, *in vivo* analysis of choroid plexuses revealed increased macrophage trafficking and a scattered presence of polarized cells upon induction of anti-CNS inflammation.

Altogether, we show that acquisition of distinct macrophage polarizations significantly alters the adhesive and migratory properties of these cells in a barrier-specific fashion. While monocytes trafficking at the level of the BBB seem to acquire their signature phenotype only following diapedesis, other anatomical interfaces can be the entry site for functionally activated monocyte-derived cells. Our study highlights the choroid plexus as a key access gateway for macrophages during neuroinflammation, and its stroma as a potential priming site for their functional polarization.

## INTRODUCTION

Central nervous system (CNS) interfaces are complex, multilayered structures hosting different types of yolk sac-derived barrier-associated macrophages^1^. While new tools have recently allowed a detailed categorization of these cells^2–4^, their functional contribution to tissue homeostasis at organ interfaces remains unclear^5^.

At the same time, CNS borders comprise anatomical specializations which limit the trafficking of pathogens and of blood-borne immune cells toward the organ parenchyma^6^. Tight barriers are present at the level of the leptomeningeal vasculature within the subarachnoid space, of the blood-cerebrospinal fluid barrier (BCSFB) within the choroid plexus, and of the blood-brain barrier (BBB) within CNS microvascular endothelial cells^7^. Altogether, access to the CNS parenchyma from the peripheral immune system appears a dynamic but tightly regulated process. Initially, passage through CNS gateways leads to cells accumulating from the BBB in the perivascular space and from the BCSFB and the leptomeningeal vessels in the cerebrospinal fluid (CSF) and the subarachnoid space^7^. These anatomical compartments are separated from the CNS parenchyma by a basement membrane and glia limitans^8^, hence constituting a second “checkpoint” mechanism to further limit contact of blood borne cells with glial and neuronal cells^9^.

However, during neuroinflammatory diseases such as Multiple Sclerosis (MS), bone marrow-derived immune cells gain access to different CNS areas. By trafficking through organ interfaces, they drastically increase barrier complexity eventually reaching the CNS parenchyma^10^, where they guide disease progression^11^. Multifocal CNS infiltration of immune cells through (compromised) CNS barriers is also a pathological hallmark of experimental autoimmune encephalomyelitis (EAE). This animal model of auto-aggressive CNS immunity is widely used to study immune cell trafficking to the CNS, despite its intrinsic limitations as a pure MS model^12^.

Within CNS lesions in both MS and EAE, monocyte-derived macrophages are the most represented infiltrating immune cells^13,14^. In the mouse system, these players arise from inflammatory and patrolling circulating monocytes, respectively characterized by high and low expression of the differentiation marker Ly6C^15^. These distinct cell types display differential expression of the chemokine receptors CCR2 and CX3CR1^16^, with Ly6C^low^CX3CR1^high^ patrolling cells required to maintain endothelial homeostasis, and inflammatory Ly6C^high^CX3CR1^low^ monocytes showing fast CCR2-mediated recruitment to inflamed tissues^17,18^. Once in the CNS, these cells coordinate the overall immune response, exerting a wide array of effector functions ranging from damaging pro-inflammatory to tissue repairing anti-inflammatory actions^19–21^.

The remarkable plasticity of monocyte-derived cells originates from the integration of numerous local signals leading to a dynamic functional modulation^22^, with pro- and anti-inflammatory cells characterized by distinct metabolic states and by differential regulation of phenotypic markers^19^. Among these, expression of the enzymes iNOS and arginase-1 is commonly used as signature marker for pro- and anti-inflammatory macrophage polarizations, respectively^23^. These two proteins utilize L-arginine as common substrate, leading to the production of cytotoxic nitric oxide and citrulline (iNOS), or urea and ornithine (arginase-1) with key downstream tissue repair properties^24^.

By visualizing expression of reporter proteins under the control of the *Nos2* and *Arg1* promoters, we have recently described the functional evolution of CNS-infiltrating macrophages in the EAE animal model^25^. Inflammatory lesion formation at a pre-clinical stage was characterized by parenchymal invasion of CCR2^+^ Ly6C^high^ iNOS^+^ monocyte-derived cells. These cells revealed an innate functional plasticity by progressively switching *in vivo* their phenotype toward an iNOS^negative^-arginase-1^+^ state throughout disease evolution. This transition was paralleled by the distinct CNS infiltration of arginase-1^+^ phagocytes that did not previously express iNOS^25^.

While no substantial iNOS/arginase-1 expression could be detected in the periphery of immunized mice, detailed CNS analysis revealed a compartmentalized distribution of distinct phenotypes within anatomical CNS interfaces. First, leptomeningeal spaces revealed a significant increase in the density of arginase-1^+^ phagocytes, as opposed to the iNOS^+^-dominated nature of parenchymal lesions; secondly, pro-inflammatory phagocytes were observed at high densities in the perivascular space and at the meningeal/parenchymal interface, with freshly-invading phagocytes acquiring iNOS expression in such transitional location^25^.

While local cues^19,22^ can contribute to the evolution of functional polarizations within the CNS, the guiding factors and the sub-anatomical CNS compartments where monocytes acquire their phenotype are not known. Furthermore, whether pro- and anti-inflammatory macrophages are preferentially recruited at specific CNS gateways remains unclear. This hinders the design of therapeutic interventions selectively blocking the invasion pathway of cytotoxic players.

In this study, we assessed the interaction of differentially polarized monocyte-derived cells with endothelial (BBB) and epithelial (BCSFB) barriers of the CNS in tissue sections and functionally *in vitro*. While several models of brain barriers exist^26^, the use of primary mouse brain barrier models allowed direct comparison with *in vivo* studies, altogether modelling how committed macrophages interact with different CNS interfaces. Here we show that, while passage of monocyte-derived cells across the BBB endothelium is strongly reduced upon functional polarization, pro- and anti-inflammatory macrophages can efficiently interact with the BCSFB following induction of CNS autoimmune inflammation. Altogether, our data prove that intraluminal monocytes at the BBB could acquire their functional specifications only during/following diapedesis through an activated endothelium, while other CNS border areas such as the choroid plexus BCSFB can be an efficient CNS access gateway for fully committed pro and anti-inflammatory monocyte-derived cells. Further studies are now needed to understand the trafficking and behavior of macrophages at other CNS interfaces such as the leptomeninges, and the potential polarizing effects of distinct CNS barrier structures on circulating monocytes and monocyte-derived cells.

## RESULTS

### Functional characterization of primary mouse brain barrier model

*In vitro* models of brain barriers are a valuable tool to study dynamics of leukocyte interaction with endothelial and epithelial cells^27^.

To investigate adhesion and transmigration of mouse monocyte-derived cells, we first isolated primary mouse brain microvascular endothelial cells (pMBMECs, to mimic the BBB^28,29^) and primary mouse choroid plexus epithelial cells (pMCPECs, to mimic the BCSFB^30^) according to well-established approaches.

pMBMECs formed a confluent monolayer, characterized by junctional localization of the tight junction proteins claudin-5 and zona occludens-1 (ZO-1) in addition to the adherens junction proteins VE-cadherin and β-catenin (**Fig. 1**). To model BBB activation occurring during neuroinflammatory conditions, pMBMECs were stimulated either with interleukin-1β (IL-1β) or with tumor necrosis factor-α (TNF-α) + interferon-γ (IFN-γ)^29,31^. Cell surface protein expression of integrin-binding molecules such as intercellular adhesion molecule-1 (ICAM-1) and vascular cell adhesion molecule-1 (VCAM-1) was upregulated upon cytokine stimulation compared to unstimulated pMBMECs (**Fig. 2a**). Additionally, increased cell surface expression of the endothelial cell adhesion molecule E-selectin was observed upon IL-1β stimulation compared to TNF-α+IFN-γ and unstimulated conditions (**Fig. 2a**). As a measure of barrier integrity, we assessed trans-endothelial electrical resistance (TEER) of the pMBMECs and observed a statistically significant decrease of TEER following cytokine stimulation compared to unstimulated conditions (**Fig. 2b**).

**Figure 1.**
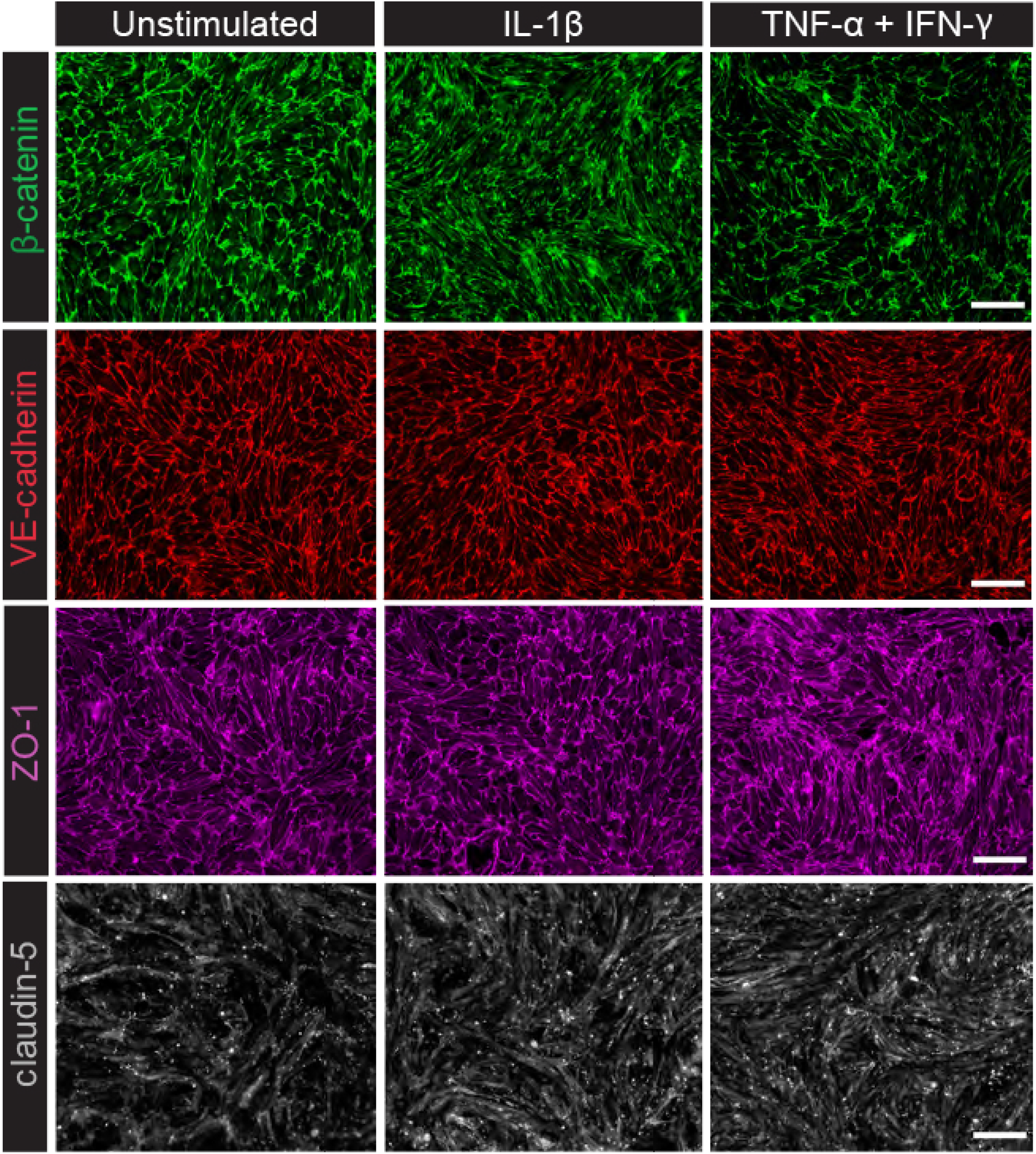
Primary Mouse Brain Microvascular Endothelial Cells (pMBMECs) characterisation. pMBMECs were isolated from *C57BL/6j* mice and grown in culture for seven days. At culture day six, pMCPECs were stimulated with 20ng/ml IL-1β or with 5ng/ml TNF-a +100IU/ml IFN-γ. Unstimulated pMBMECs were used as control. Shown are immunofluorescence stainings of adherens junction molecules β-catenin and VE-cadherin and tight junction molecules ZO-1 and claudin-5, scale bar 50μm. Pictures are representative of three independent experiments.

**Figure 2.**
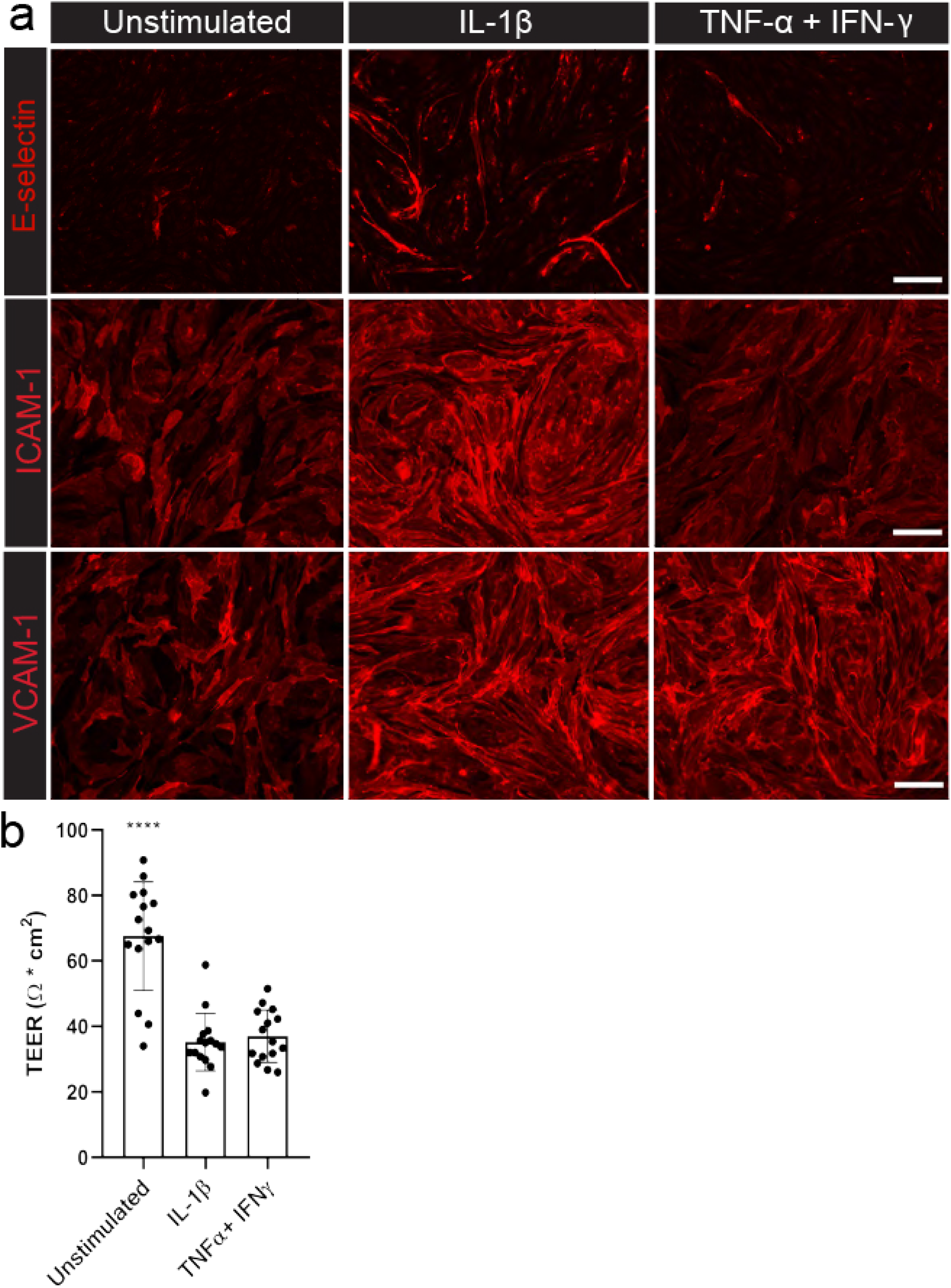
Expression of adhesion molecules and pMBMEC permeability. **a)** Surface expression of E-selectin, ICAM-1 and VCAM-1 on unstimulated and IL-1β or TNF-α+IFN-γ stimulated pMBMECs at culture day seven. Pictures are representative of three independent experiments, 50μm scale bar. **b)** Tightness of pMBMECs cultured on 5μm pore size filters was determined by TEER (Ω*cm^2^). TEER of stimulated pMBMECs is significantly lower compared to unstimulated endothelium [One-way ANOVA, F(2,42) = 35.86, *p* < 0.0001; IL-1β pMBMECs: Mean = 35.20, STDev = 8.73; TNF-α+IFN-γ pMBMECs: Mean = 36.98, STDev = 7.97; Unstimulated pMBMECs: Mean = 67.56, STDev = 16.59].

To increase comparability between different experimental setups, pMCPECs as *in vitro* model for the BCSFB were isolated from the same mice that yielded pMBMECs. In accordance to previous observations, pMCPECs reached confluence following six days in culture. Epithelial monolayers displayed mature BSCFB characteristics as shown by the localization of junctional molecules of E-cadherin, β-catenin, junctional adhesion molecule-A (JAM-A) and claudin-1 (**Fig. 3**). At culture day seven, pMCPECs were stimulated either with TNF-α or IFN-γ to mimic local inflammation or were left unstimulated^30^. IFN-γ stimulation led to a significant upregulation of ICAM-1 compared to stimulation with TNF-α or unstimulated conditions (**Fig. 4a**). Barrier permeability was determined by trans-epithelial electrical resistance (TEER) measurement of cellular monolayers cultured on 5μm pore filters. No statistically significant differences in permeability were observed between stimulated and unstimulated epithelial cells (**Fig. 4b**).

**Figure 3.**
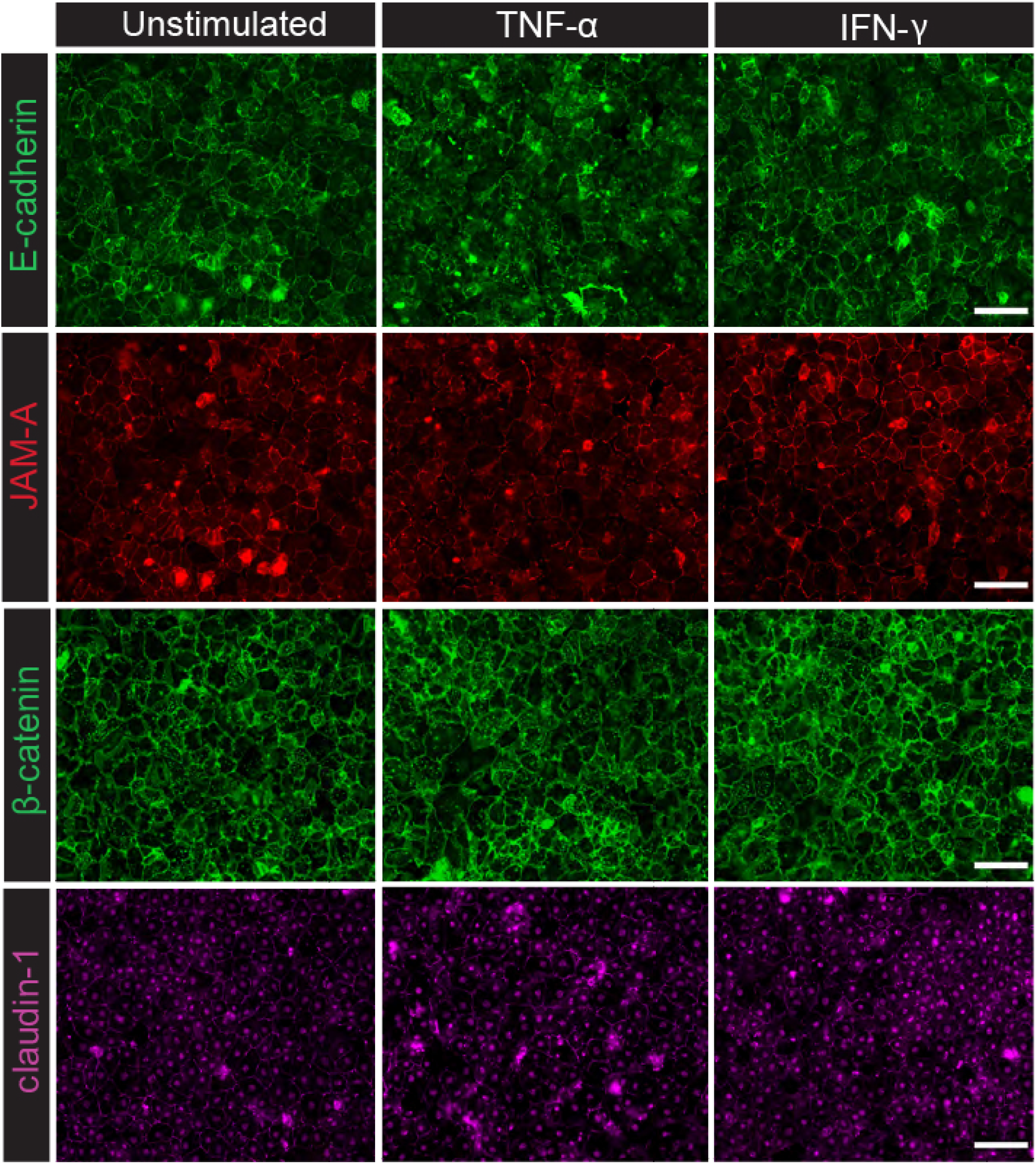
Primary Mouse Choroid Plexus Epithelial Cells (pMCPECs) characterisation. pMCPECs were isolated from lateral, 3^rd^ and 4^th^ brain ventricles of *C57BL/6j* mice and grown in culture for seven days. At culture day six, pMCPECs were stimulated with 10ng/ml TNF-α or with 100IU/ml IFN-γ. Unstimulated pMCPECs were used as control condition. Shown are immunofluorescence stainings of adherens junction molecules E-cadherin, β-catenin and tight junction molecules junctional adhesion molecule-A (JAM-A) and claudin-1 expressed on unstimulated and TNF-α or IFN-γ stimulated pMCPECs at culture day seven. Pictures are representative of two independent experiments, scale bar 50μm.

**Figure 4.**
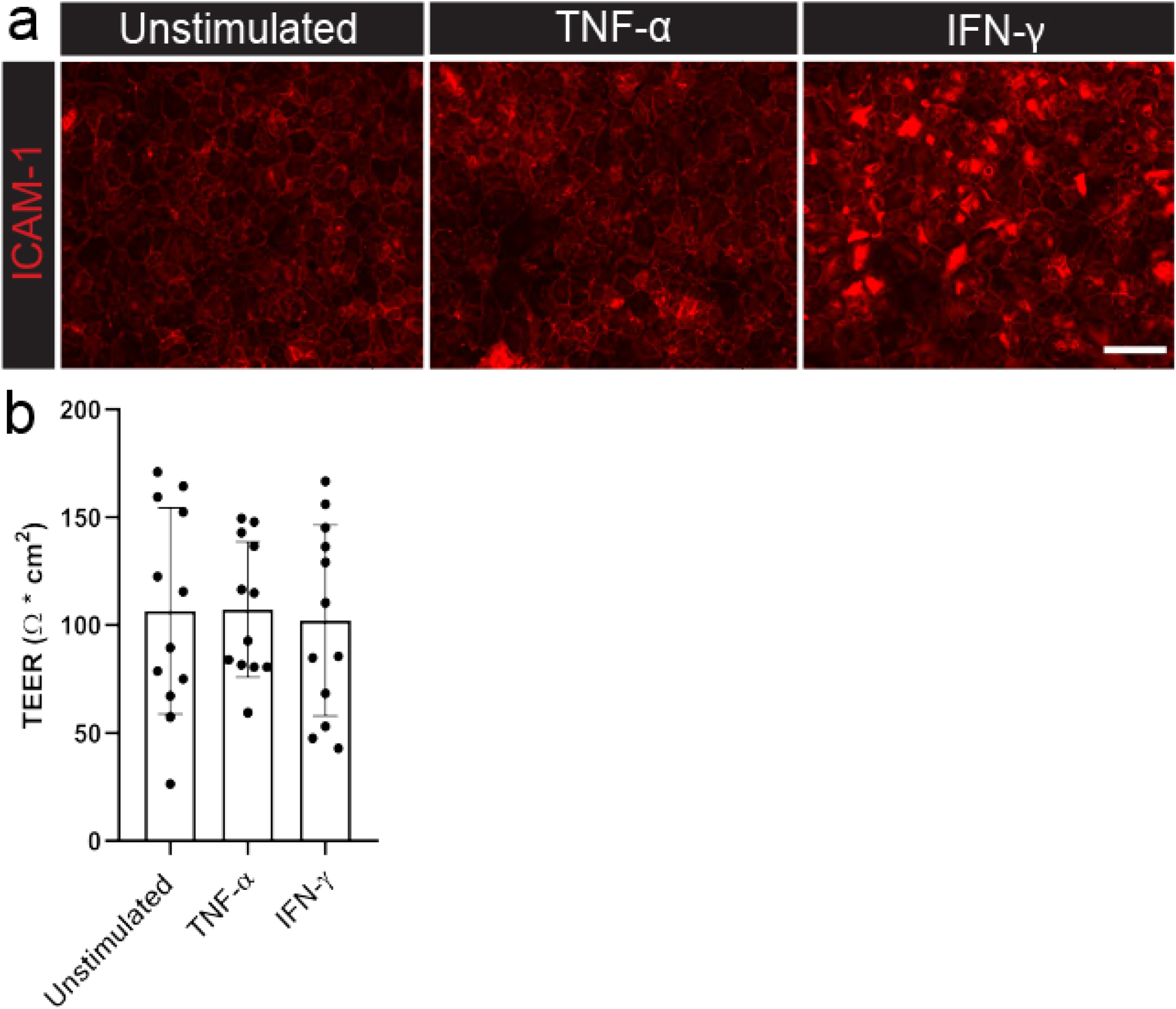
Expression of ICAM-1 and pMCPEC permeability. **a)** Surface expression of ICAM-1 on unstimulated and TNF-α or IFN-γ stimulated pMCPECs at culture day seven. Pictures are representative of two independent experiments, 50μm scale bar. **b)** Tightness of pMCPECs cultured on 5μm pore size filters was determined by TEER (Ω*cm^2^). No statistically significant differences were detected between unstimulated and TNF-α or IFN-γ stimulated pMCPECs at culture day seven by one-way ANOVA.

### *In vitro* polarization of bone marrow-derived macrophages to model pro- and anti-inflammatory *in vivo* phenotypes

To model primary monocyte-derived cells trafficking to and through distinct brain barriers, we made use of bone marrow-derived macrophage cultures (BMDMs). Cells were differentiated *in vitro* and kept unstimulated (M^unpolarized^) or polarized towards a pro-inflammatory (M^LPS+IFNy^) or anti-inflammatory phenotype (M^IL-4+IL-13^) to mimic the macrophage activation spectrum observed *in vivo* in inflammatory models^25^. Given the heterogeneous and confusing terminology still affecting the reporting of macrophage investigations, we implemented a stimulus-based nomenclature (M^unpolarized^, M^LPS+IFNy^, M^IL-4+IL-13^) according to suggestions by experts in the field^32^. As expected, M^LPS+IFNy^ cells displayed a significant upregulation of pro-inflammatory genes such as *Nos2, Cd38, Fpr2, Cd86, Stat1, Gpr18* (**Fig. 5a**) whereas M^IL-4+IL-13^ cells showed increase in expression of signature genes *Retnla, Arg1, Ym1, Egr2* and *Mrc1* (**Fig. 5b**).

**Figure 5.**
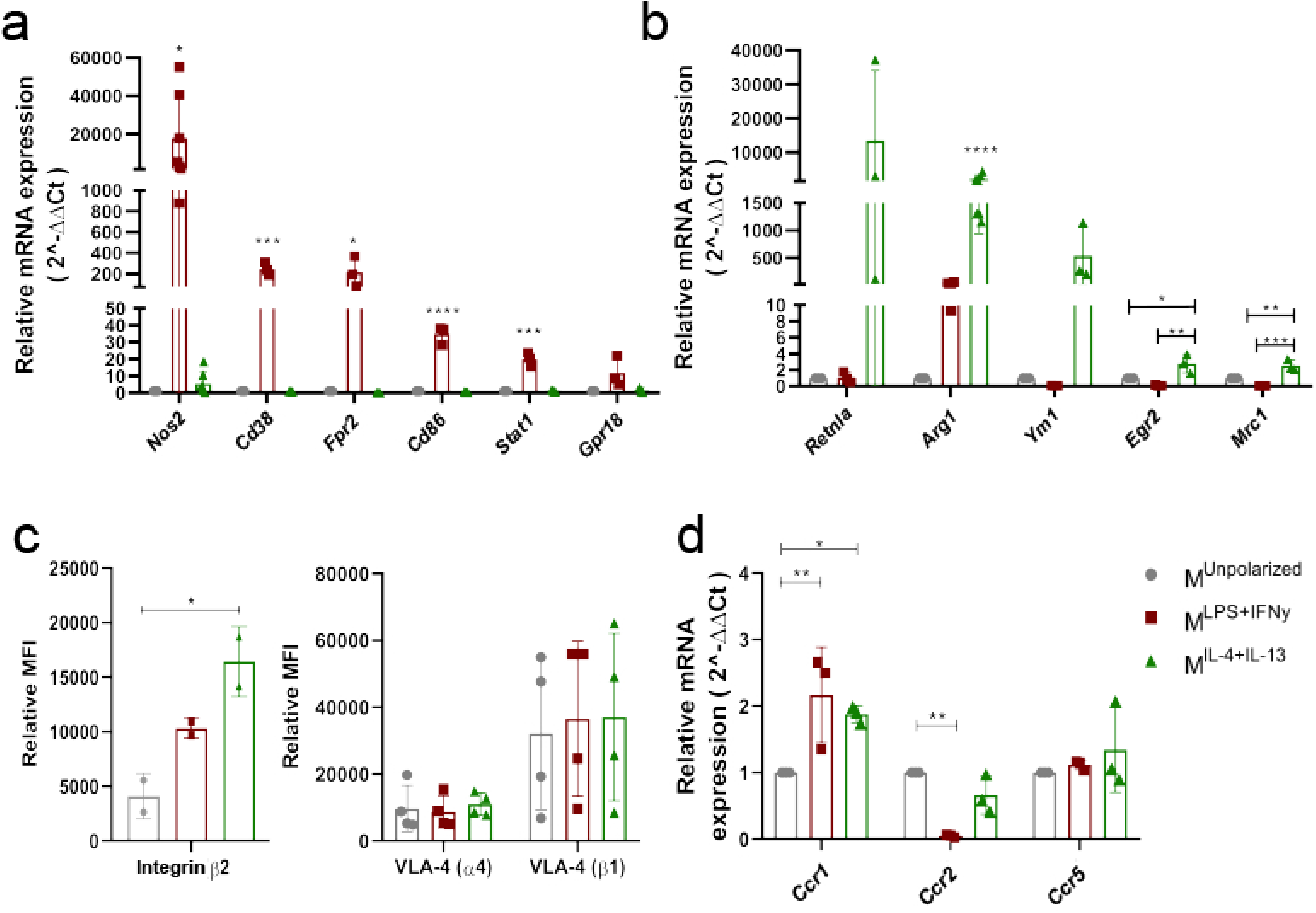
Characterization of BMDM cultures. BMDMs were stimulated with 100ng/ml LPS + 10ng/ml IFN-γ (M^LPS+IFN-γ^) or with 10ng/ml IL-4 + 10ng IL-13 (M^IL-4+Il-13^) at culture day seven. M^unpolarized^ cells were left unstimulated. **a,b**) Gene expression was assessed by RTqPCR following mRNA extraction. Data is normalized using *Hprt* as reference gene and presented as fold increase relative to M^unpolarized^ condition (2^−ΔΔCt^, see Methods). **a**) Relative mRNA expression of pro-inflammatory genes *Nos2, Fpr2, Gpr18, Cd38, Cd86, Stat1* and of **b**) anti-inflammatory genes *Retnla, Arg1, Mrc1, Ym1* and *Egr2* is shown. Displayed are means ± standard deviation obtained from technical triplicates, 3 independent experiments. M^LPS+IFN-γ^ compared to M^IL-4+Il-13^ and to M^unpolarized^ cells have a significantly higher expression of *Nos2* [one-way ANOVA, F(2,18) = 4.677, *p* = 0.023]*, Fpr2* [one-way ANOVA, F(2,6) = 6.587, *p* = 0.031]*, Cd38* [one-way ANOVA, F(2,6) = 46.46, *p* =0.0002]*, Cd86* [one-way ANOVA, F(2,6) = 114.5, *p* < 0.0001]*, Stat1* [one-way ANOVA, F(2,6) = 63.48, *p* < 0.0001]. Compared to M^unpolarized^ and M^LPS+IFN-γ^ macrophages, M^IL-4+IL-13^ cells display significantly higher expression of *Arg1* [one-way ANOVA, F(2,18) = 22.13, *p* < 0.0001], *Egr2* [one-way ANOVA, F(2,6) = 12.68, *p* = 0.007] and *Mrc1* [one-way ANOVA, F(2,6) = 30.89, *p* = 0.007]. **c**) Surface expression of β2, α4 and β1 integrin molecules detected on M^unpolarized^, M^LPS+IFN-γ^ and M^IL-4+Il-13^ macrophages by flow cytometry. Displayed are relative mean fluorescence intensity (MFI) values obtained by subtracting the MFI of stained samples from the MFI of isotype control-stained samples; data presented as mean ± standard deviations from 2 independent experiments for β2 integrin and from 4 independent experiments for α4 and β1 integrins. M^IL-4+Il-13^ display a significantly increased β2 integrin expression compared to M^LPS+IFN-γ^ and M^unpolarized^ cells [one-way ANOVA, F(2,3) = 14.92, *p* = 0.027]. **d**) Relative mRNA expression of chemokine receptors 1, 2 and 5 (*Ccr1*, *Ccr2*, *Ccr5*) in M^unpolarized^, M^LPS+IFN-γ^, M^IL-4+Il-13^ cells after 48h cytokine stimulation. Upon polarization, astatistically significant increase in *Ccr1* expression was observed in M^LPS+IFN-γ^ (p = 0.001) and M^IL-4+Il-13^ (p = 0.013) compared to unstimulated cells. M^LPS+IFN-γ^ displayed significantly downregulated *Ccr2* expression compared to M^unpolarized^ cells (p = 0.007), one-way ANOVA. Shown is mean ± standard deviation from 3 independent experiments.

Leukocyte extravasation is a dynamic multistep cellular interaction process which sequentially involves rolling, cell arrest and firm adhesion, slow crawling on the endothelium and eventually paracellular or transcellular diapedesis across the endothelial wall^33^. The fine dynamics of this process and the molecules involved vary depending on the anatomical and inflammatory context^34^, but typically rely on the expression and activation of key adhesion molecules such as selectins, leukocyte integrins and adhesion molecules of the immunoglobulin superfamily^35,36^. Flow cytometric analysis of integrin expression in BMDMs revealed high basal expression of β2, α4 and β1 integrin subunits, with β2 integrins significantly more expressed in M^IL-4+IL-13^ cells^37^ (**Fig. 5c**).

Integrin activation prior to luminal crawling is also mediated by signaling through chemokine receptors^33^ in a phenotype-dependent manner^38^. Furthermore, chemokine stimulation can also influence macrophage polarization in pathological contexts^39^. We thus investigated expression of the main chemokine receptors *Ccr1*, *Ccr2* and *Ccr5* on BMDMs at the RNA level. While *Ccr5* levels were comparable in M^unpolarized^, M^LPS+IFNy^ and M^IL-4+IL-13^ cells, we could observe a significant increase of *Ccr1* in M^LPS+IFNy^ and M^IL-4+IL-13^, and a substantial decrease of *Ccr2* expression in M^LPS+IFNy^ cells compared to M^IL-4+IL-13^ and M^unpolarized^ cells (**Fig. 5d**).

Altogether, acquisition of distinct functional states significantly affects expression of key molecules involved in cell dynamics. Given the recognized role of CCL2 signaling in macrophage-driven clinical development of EAE^40^, the observed decrease of *Ccr2* expression in M^LPS+IFNy^ BMDMs might specifically affect the trafficking properties of these model cells.

### BMDM polarization decreases adhesion to the BBB and migration *in vitro*

Sensing of inflammation by circulating monocytes initiates in the vascular lumen following interaction with activated endothelial cells^33^. The latter participate in the recruitment and activation of leukocytes by secreting several chemokine and inflammatory cytokines^41^, altogether exerting a complex net effect on circulating immune cells^42^. Whether monocytes can acquire a signature functional phenotype once in contact with endothelial cells, and whether this polarization affects their adhesive and extravasating properties is however not known. While *in vivo* imaging offers an invaluable tool to describe cell dynamics at the (superficial) meningeal and subpial level^25^, the properties of parenchymal BBB and intraventricular BCSFB can be recapitulated *in vitro* using specific primary models.

To study the migration of differentially polarized monocyte-derived cells across pMBMECs as a model for the BBB, we used a transmigration system in which BMDMs were stained with a live cell tracker dye (CMFDA), incubated on top of pMBMECs and allowed to migrate across these monolayers for 8h. In this assay, while an irrelevant minority of BMDMs could be detected suspended in the lower well medium by flow cytometry (data not shown), most macrophages adhered to the luminal endothelial surface (**Fig. 6a**) or migrated to the endothelial abluminal side, remaining however in contact with the filter (**Fig. 6b**).

**Figure 6.**
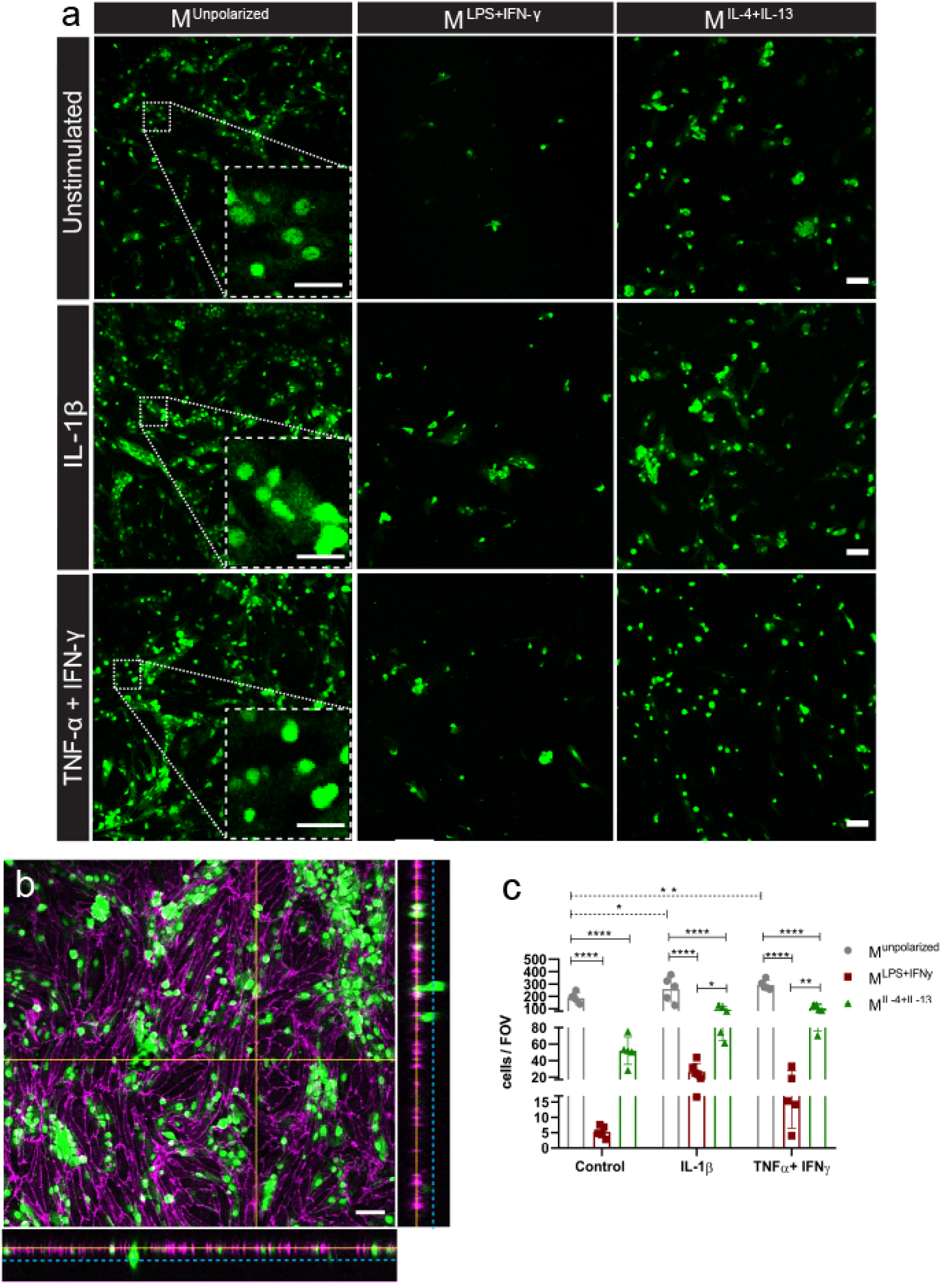
BMDM adhesion to pMBMECs *in vitro*. **a)** Representative images illustrating CMFDA^+^ M^unpolarized^, M^LPS+IFN-γ^ and M^IL-4+Il-13^ cells attached on the luminal side of unstimulated, IL-1β or TNF-α+IFN-γ stimulated pMBMECs following 8h incubation. Scale bar, 50 μm; scale bar of magnified representative inlets, 25 μm. **b)** Representative confocal z-stack of M^unpolarized^ cells on a filter containing unstimulated pMBMECs. 90° X and Y projections (created with Fiji) show relative location of macrophages compared to endothelial cells; yellow line indicates macrophages on the luminal side of pMBMECs, dotted cyan line indicates macrophages that migrated through the pMBMECs on the abluminal side. Immunostaining with anti-ZO-1 antibody, purple; CMFDA^+^ macrophages, green. Scale bar, 50 μm. **c)** Number of M^unpolarized^, M^LPS+IFN-γ^ and M^IL-4+Il-13^ cells attached on the luminal side of unstimulated and IL-1β or TNF-α+IFN-γ stimulated pMBMECs following 8h incubation. Data points represent mean number of cells per filter, with five fields of view (FOV) analysed, for a total of three individual experiments, 4 filters per condition. In all conditions, M^unpolarized^ compared to M^LPS+IFN-γ^ and M^IL-4+Il-13^, p > 0.0001. Compared to M^LPS+IFN^-γ, M^IL-4+Il-13^ cells adhere significantly more to IL-1β (p = 0.033) and TNF-α+IFN-γ stimulated pMBMECs (p = 0.005). M^unpolarized^ cells adhere in higher numbers to both IL-1β (p = 0.018) and TNF-α+IFN-γ pMBMECs (p = 0.0006) compared to control conditions. Two-way ANOVA with Tukey’s multiple comparisons test, *p < 0.05, **p < 0.01, ***p < 0.001.

Using confocal microscopy, we could observe that M^unpolarized^ cells adhered in a significantly higher number to the apical side of pMBMECs compared to polarized M^LPS+IFNy^ and M^IL-4+IL-13^ macrophages in all experimental conditions (**Fig. 6a,c**). Moreover, activation of pMBMECs with the pro-inflammatory cytokines TNF-α+IFN-γ or IL-1β further increased M^unpolarized^ attachment to endothelial cells (**Fig. 6c**).

When assessing the number of macrophages on the abluminal pMBMEC side, we observed accordingly a significantly higher presence of transmigrated M^unpolarized^ compared to M^LPS+IFNy^ and M^IL-4+IL-13^ cells (**Fig. 7a,b**). Passage of M^unpolarized^ and M^IL-4+IL-13^ cells, but not of M^LPS+IFNy^ cells, was increased following pMBMEC stimulation with TNF-α+IFN-γ or IL-1β (**Fig. 7b**). Altogether, pro-inflammatory skewing virtually abolished BMDM attachment to the BBB endothelium (**Fig. 6c and 7b**). Interestingly however, we could observe that the intrinsic extravasating efficiency of M^LPS+IFNy^ cells was similar to M^unpolarized^ cells in all culture conditions, but significantly inferior to the diapedesis rate of M^IL4+IL13^ when incubated with IL-1β-stimulated pMBMECs (**Fig. 7c)**. In conclusion, we could show in our *in vitro* system that M^unpolarized^ cells efficiently adhere and migrate across unstimulated and activated pMBMECs modeling the BBB; however, upon anti-inflammatory and even more drastically following pro-inflammatory polarization, BMDM interaction with pMBMECs was significantly reduced.

**Figure 7.**
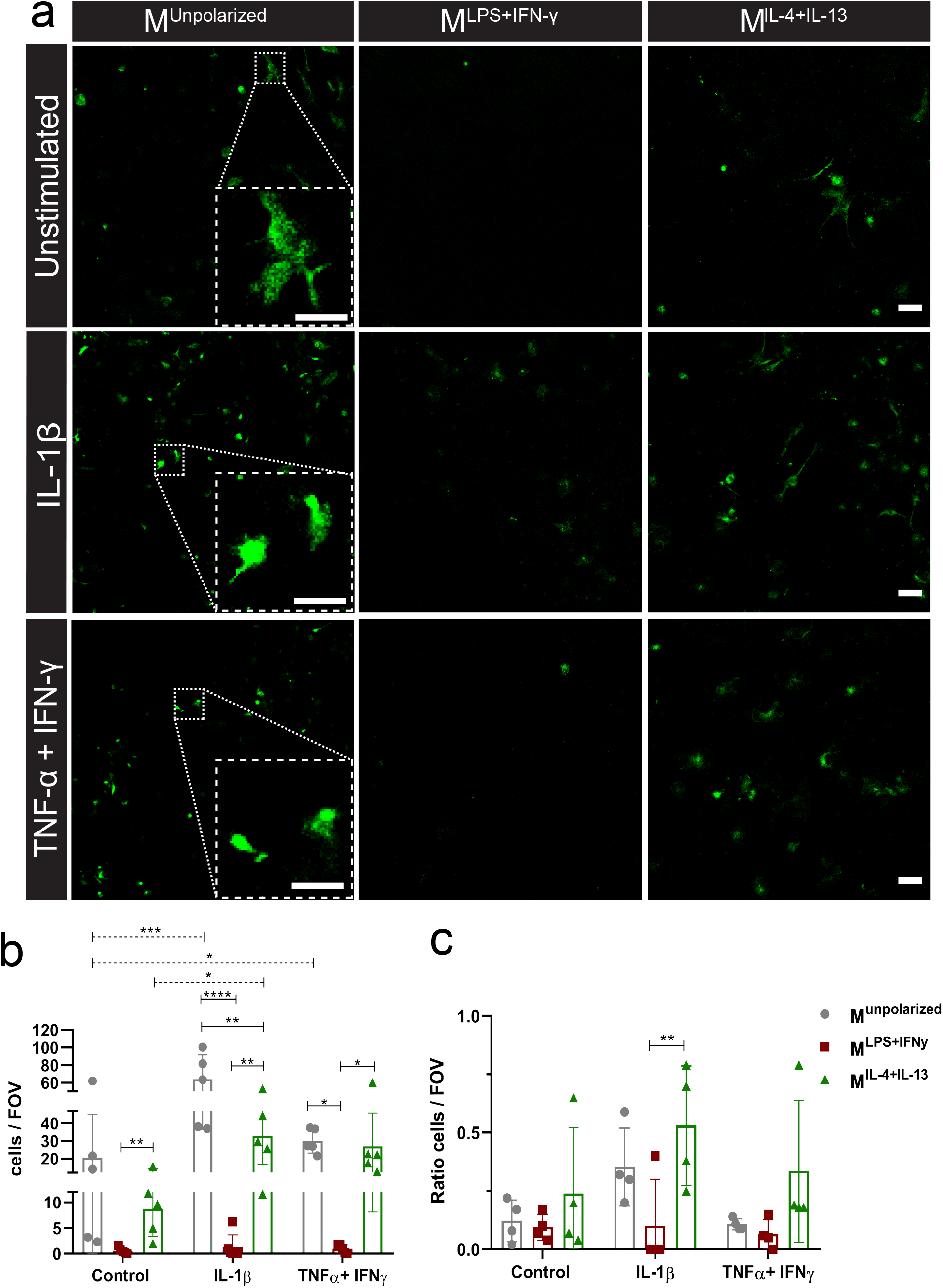
BMDM migration across pMBMECs *in vitro*. **a)** Representative images of CMFDA^+^ M^unpolarized^, M^LPS+IFN-γ^ and M^IL-4+Il-13^ cells migrated to the abluminal side of unstimulated, IL-1β and TNF-α+IFN-γ stimulated pMBMECs following 8h incubation. Scale bar, 50 μm; scale bar of magnified inlets, 25 μm. **b)** Number of BMDMs on the abluminal side of unstimulated and IL-1β or TNF-α+IFN-γ stimulated pMBMECs after 8h incubation. Data points represent mean number of cells per filter (five fields of view per sample, FOV). Three individual experiments performed, 4 filters per condition. A significantly higher number of M^unpolarized^ cells crossed IL-1β stimulated endothelium compared to both M^LPS+IFN-γ^ (p < 0.0001) and M^IL-4+Il-13^ (p = 0.0057) cells. M^unpolarized^ cells show increased migration across TNF-α+IFN-γ pMBMECs compared to M^LPS+IFN-γ^ cells (p < 0.011). M^IL-4+Il-13^ cells migrated in higher numbers across IL-1β (p = 0.004) and TNF-α+IFN-γ pMBMECs (p = 0.024) compared to M^LPS+IFN-γ^ macrophages. M^unpolarized^ cells migrate more effectively through IL-1β (p =0.0001) and TNF-α+IFN-γ pMBMECs (p = 0.003) compared to unstimulated pMBMECs; M^IL-4+Il-13^ migrate in higher numbers across IL-1β stimulated pMBMECs than across unstimulated endothelium (p = 0.04). Two-way ANOVA with Tukey’s multiple comparisons test, *p < 0.05, **p < 0.01, ***p < 0.001. **c)** Bar graph illustrates the ratio of M^unpolarized^, M^LPS+IFN-γ^ and M^IL-4+Il-13^ macrophages attached on the luminal side of pMBMECs compared to the number of macrophages on the abluminal side of pMBMECs following 8h incubation. Shown are the average migrated/attached cells ratios per filter, obtained by quantifying the number of cells from five different fields of view (FOV). Three individual experiments were performed, 4 filters analyzed per condition. M^IL-4+Il-13^ displayed a higher migration ratio on IL-1β pMBMECs compared to M^LPS+IFN-γ^ cells (p = 0.009). Two-way ANOVA with Tukey’s multiple comparisons test, *p < 0.05, **p < 0.01, ***p < 0.001.

### BMDM polarization abolishes adhesion to the BBB *in vitro* under shear flow

Physiological blood flow plays a critical role in immune cell migration across the BBB, with shear forces within the vascular lumen allowing tighter interactions of leukocytes with endothelial surfaces^43^. To mimic lumen fluid dynamics experienced by migrating monocytes, we used a previously established live cell imaging approach^44^ and allowed BMDMs to interact with pMBMEC monolayers for 30 minutes under flow conditions (1.5 dyn/cm^2^). Similarly to what we observed in the absence of shear flow (**Fig. 6**), both M^LPS+IFNy^ and M^IL-4+IL-13^ cells showed significantly reduced adhesion to pMBMECs compared to M^unpolarized^ macrophages (**Fig. 8a,b**). Stimulation of endothelial cells with inflammatory cytokines further increased adhesion of M^unpolarized^ cells under shear flow, albeit not significantly (**Fig. 8b**). Notably, while M^LPS+IFNy^ could adhere in low numbers only to IL-1β-activated pMBMECs, M^IL-4+IL-13^ cells did not interact with pMBMECs in any experimental condition (**Fig. 8a,b**). Following adhesion, M^unpolarized^ cells displayed highly dynamic interactions with endothelial cells, characterized by continuous probing, luminal crawling, and partial or full diapedesis across the pMBMEC monolayer (**Fig. 8c**). This behavioral pattern did not differ however between unstimulated and stimulated pMBMECs (**Fig. 8c**).

**Figure 8.**
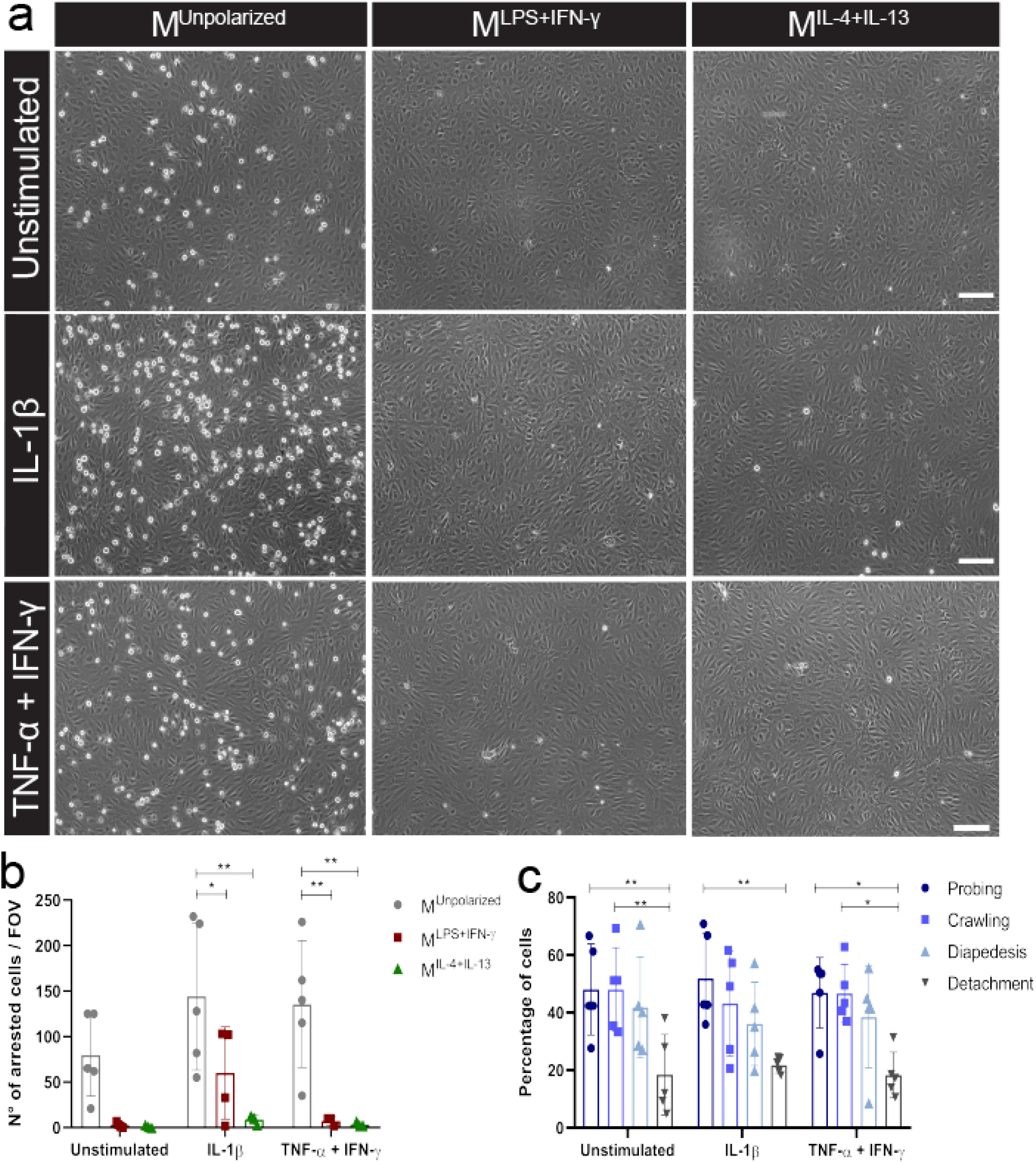
BMDM migration across pMBMECs under flow conditions. **a)** Representative images illustrating M^unpolarized^, M^LPS+IFN-γ^ and M^IL-4+Il-13^ cells which adhered on unstimulated, IL-1β and TNF-α+IFN-γ stimulated pMBMECs monolayer under physiological flow (1.5 dyn/cm^2^); scale bar 100μm. **b)** Density of arrested BMDMs on stimulated and unstimulated pMBMECs per field of view (FOV). M^unpolarized^ cells displayed superior abilities to adhere to both IL-1β (M^unpolarized^ vs M^LPS+IFN-γ^ p = 0.041; M^unpolarized^ vs M^IL-4+Il-13^ p = 0.002) and TNF-α+IFN-γ stimulated endothelial cells (M^unpolarized^ vs M^LPS+IFN-γ^ p = 0.004; M^unpolarized^ vs M^IL-4+Il-13^ p = 0.003). Displayed are the results from 4 independent experiments. **c)** Migratory behaviour of M^unpolarized^ macrophages on stimulated and unstimulated pMBMECs under physiological flow conditions, during 25 minutes live cell imaging. The majority of M^unpolarized^ cells adopted a probing and crawling behaviour following attachment to pMBMECs, whereas fewer cells fully crossed the monolayer. No significant differences were observed between conditions. A significantly lower number of cells detached from the endothelium following attachment, compared to the number of cells that crawled or probed the monolayer (unstimulated pMBMECs: probing vs detachment p = 0.009, crawling vs detachment p = 0.009; IL-1β pMBMECs: probing vs detachment p = 0.008; TNF-α+IFN-γ pMBMECs: probing vs detachment p = 0.013, crawling vs detachment p = 0.014). Statistical analysis performed by two-way ANOVA, followed by Tukey’s multiple comparison test, *p < 0.05; **p < 0.01; ***p < 0.001.

Taken together, monocyte-derived cells under shear flow show functional adhesion and transmigration properties on brain microvascular endothelial cells both under homeostatic and inflammatory conditions. Notably, acquisition of a pro- or anti-inflammatory phenotype drastically decreases these interactive properties, thus suggesting that infiltrating macrophages at the level of the BBB can become functionally polarized only during/following diapedesis through an activated endothelial wall.

### Polarized BMDMs efficiently interact with the BCSFB epithelium *in vitro*

The BCSFB hosts different types of monocyte-derived cells^2^ and constitutes a selective invading gateway for blood-borne immune cells during homeostasis and inflammation^7,45^. Accordingly, epithelial cells of the choroid plexuses express adhesion molecules and chemokines involved in cell migration^46^. Whether the BCSFB can be an entry gateway for functionally polarized monocyte-derived cells during auto-aggressive CNS conditions remains however unclear.

To mimic macrophage adhesion and migration at this CNS gateway, we incubated CMFDA-stained M^unpolarized^, M^LPS+IFNy^ and M^IL-4+IL-13^ cells with pMCPECs cultured on inverted filters, thus simulating the physiological orientation of epithelial cells within the organ^30^. After 8 hours, we assessed by confocal microscopy the number of CMFDA^+^ monocyte-derived cells attached to the filter, on the basolateral side of pMCPECs (**Fig. 9a,b**). M^unpolarized^ and M^IL-4+IL-13^ cells adhered in significantly higher numbers to the basolateral side of unstimulated pMCPECs compared to M^LPS+IFNy^ cells (**Fig. 9a,c**). Conversely, all M^unpolarized^, M^LPS+IFNy^ and M^IL-4+IL-13^ cells could efficiently attach to the basolateral side of cytokine-stimulated pMCPECs (**Fig. 9a,c**). While BMDMs placed on control filters without pMCPECs did not migrate from the upper part of the insert during the incubation period (data not shown), in the presence of epithelial cells M^unpolarized^, M^LPS+IFNy^ and M^IL-4+IL-13^ cells could move across the filter and the cell monolayer (**Fig. 9b**). When analyzing the number of BMDMs migrating toward the apical side of pMCPECs, we could observe a comparable absolute number of M^LPS+IFNy^, M^IL-4+IL-13^ and M^Unpolarized^ cells, with no major differences between distinct pMCPEC stimulation settings (**Fig. 10a,b**). Interestingly, when assessing the ratio of macrophages crossing the pMCPEC monolayer compared to the number of cells attached on the basolateral side of the filter, relatively more M^LPS+IFNy^ cells could efficiently migrate toward the apical pMCPEC side (**Fig. 10c**).

**Figure 9.**
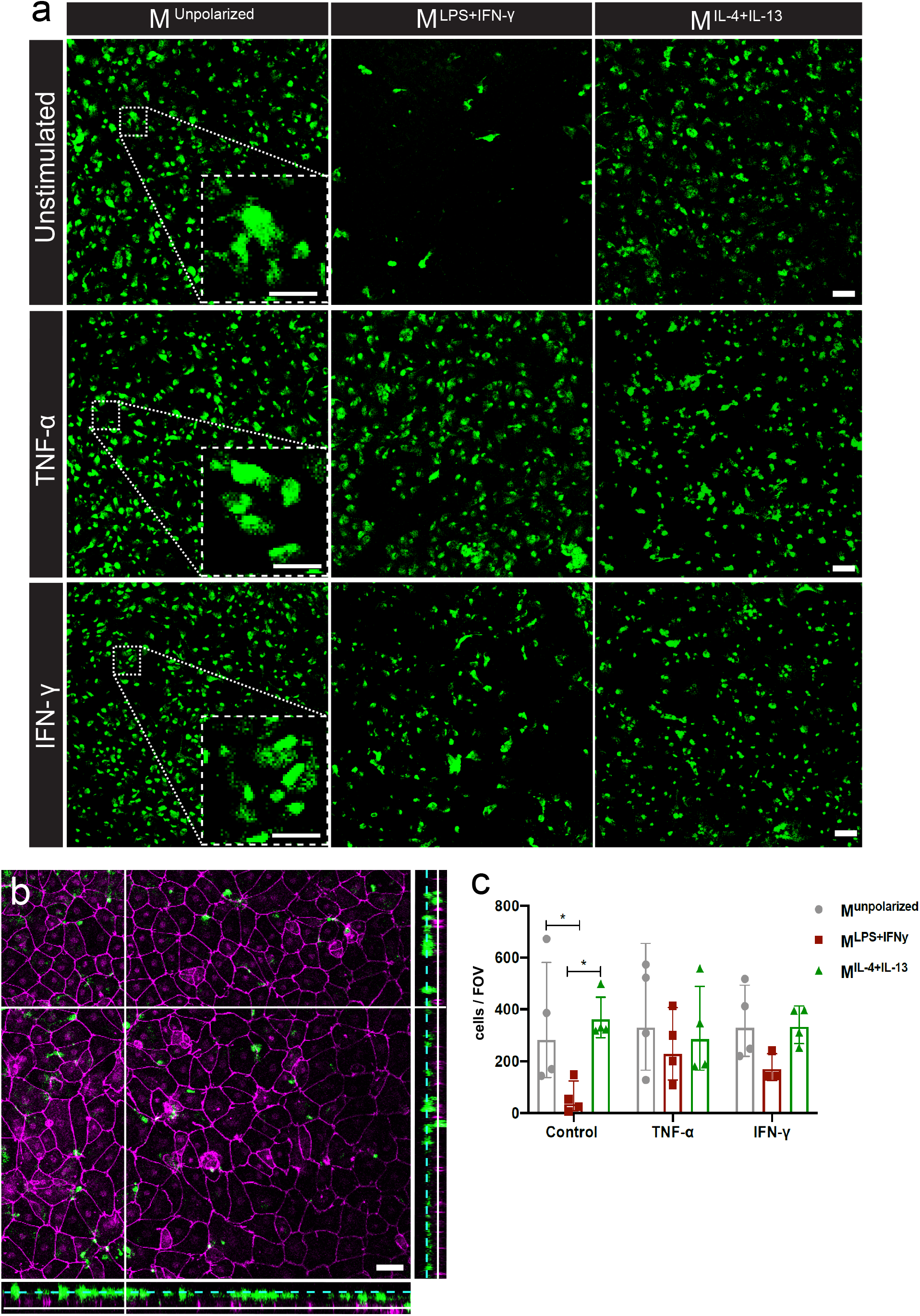
BMDM adhesion to pMCPECs *in vitro*. **a)** Representative images of CMFDA^+^ M^unpolarized^, M^LPS+IFN-γ^ and M^IL-4+Il-13^ cells attached on filters on the basolateral side of unstimulated, TNF-α or IFN-γ stimulated pMCPECs. Scale bar, 50 μm; scale bar of magnified representative inlets, 25 μm. **b)** Representative confocal z-stack of M^unpolarized^ cells on a filter containing unstimulated pMCPECs. 90° X and Y projections (created with Fiji) show relative location of macrophages compared to epithelial cells; dotted cyan line indicates macrophages attached on the basolateral side of pMCPEC-containing filters, white line indicates macrophages in between and on the apical side of pMBMECs. Immunostaining with anti-ZO-1 antibody, purple; CMFDA^+^ macrophages, green. Scale bar, 50 μm. **c)** Numbers of M^unpolarized^, M^LPS+IFN-γ^, M^IL-4+Il-13^ adhering on the filter on the basolateral side of pMCPECs following 8h incubation. M^unpolarized^ and M^IL-4+Il-13^ display superior abilities to adhere to unstimulated pMCPECs (M^unpolarized^ *vs* M^LPS+IFN-γ^ p = 0.03; M^IL-4+Il-13^ *vs* M^LPS+IFN-γ^ p = 0.017). Shown are average numbers of attached cells per filter, from five different fields of view (FOV) per filter. Four individual experiments performed, 4 filters analyzed per condition. Two-way ANOVA with Tukey’s multiple comparisons test, *p < 0.05, **p < 0.01, ***p < 0.001.

**Figure 10.**
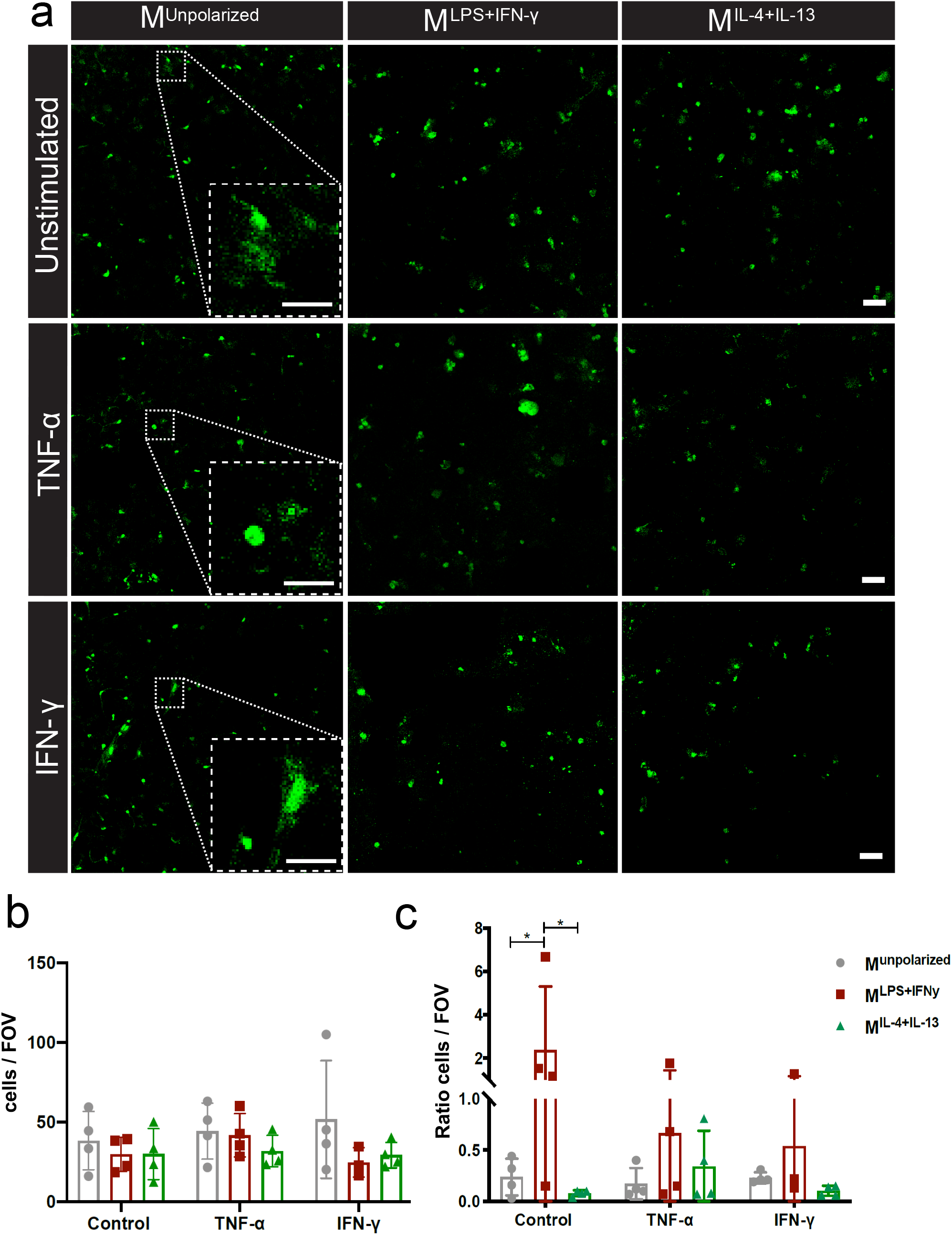
BMDM migration through pMCPECs *in vitro*. **a)** Representative images of CMFDA^+^ M^unpolarized^, M^LPS+IFN-γ^ and M^IL-4+Il-13^ cells that migrated towards the apical side of unstimulated and TNF-α or IFN-γ stimulated pMCPECs. Scale bar, 50 μm; scale bar of magnified representative inlets, 25 μm. **b)** Number of polarized and unpolarized macrophages that migrated toward the apical side of stimulated and unstimulated pMCPECs. Displayed are average numbers of migrated cells per filter, average obtained by quantifying the number of cells from five different fields of view (FOV). Four individual experiments were performed, 4 filters were analyzed per condition. **c)** Ratio between the number of BMDMs attached on the basolateral side of pMCPEC-containing filters and the number of macrophages which migrated toward the apical side of pMCPECs. M^LPS+IFN-γ^ display relative superior migration rates across unstimulated epithelium compared to M^unpolarized^ cells (p = 0.021) and M^IL-4+Il-13^ macrophages (p = 0.013). Shown are average migrated/attached cells ratios per filter, from five different fields of view (FOV) per filter. Four individual experiments were performed, 4 filters analyzed per condition. Two-way ANOVA with Tukey’s multiple comparisons test, *p < 0.05.

Taken together, these observations show that both, polarized and unpolarized macrophages can efficiently interact with the BCSFB of the choroid plexus in the absence or presence of inflammatory conditions *in vitro*.

### *In vivo* monocyte/macrophage trafficking via the choroid plexus

As our *in vitro* data suggest that the BCSFB might be an important CNS entry site for functionally committed macrophages, we used an active EAE animal model and analyzed the *in vivo* recruitment and the phenotype of monocyte-derived cells in the choroid plexus. Following EAE induction, mice developed a typical progressive pathology characterized by a preclinical loss of weight and a clinical monophasic disease evolution eventually resolving into a remission phase^25^.

Firstly, we visualized the density of infiltrating and resident monocyte-derived cells in choroid plexus-containing brain sections isolated from *Cx3CR1-GFP × CCR2-RFP* mice^17^ upon EAE induction. In this transgenic model, differential reporter expression allows distinction of CCR2^+^ blood borne macrophages from long-lived CCR2^negative^CX3CR1^high^ resident cells. In healthy animals, confocal analysis of thick choroid plexus sections revealed a minor presence of CCR2^+^ macrophages deriving from the peripheral compartment^1^, and several ramified CX3CR1^+^ resident macrophages populating the organ stroma (**Fig. 11a**, upper panel). Upon EAE immunization, we could observe a relative increase in CCR2^negative^CX3CR1^high^ cells (especially at the symptomatic peak of disease, **Fig. 11a**, lower panel), and a substantial increase in CCR2^+^ and CCR2^+^CX3CR1^low^ infiltrating cells, starting from the preclinical phase of disease (**Fig. 11a,b**). Even though not statistically significant, this augmented macrophage density suggests local proliferation of macrophages and/or enhanced recruitment of circulating CCR2^+^ monocyte-derived cells through the choroid plexus upon induction of anti-CNS immunity.

**Figure 11.**
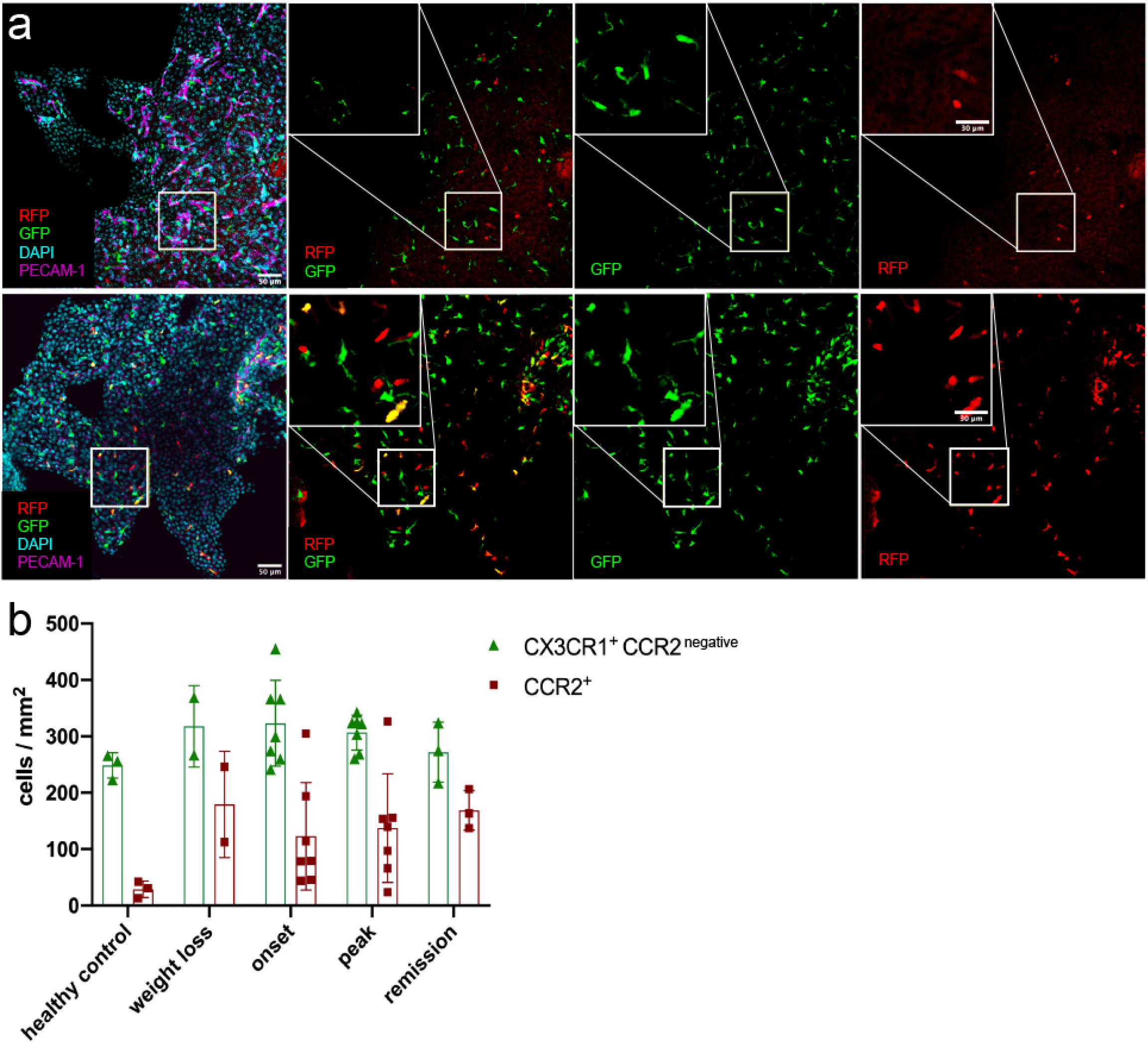
Macrophage density in the choroid plexuses of *CX3CR1-GFP × CCR2-RFP* mice during EAE. **a)** Representative confocal images of the choroid plexus from the third ventricle of a healthy *CX3CR1-GFP × CCR2-RFP* mouse (upper panel) and from a *CX3CR1-GFP × CCR2-RFP* mouse induced with EAE (weight loss stage, lower panel). RFP^+^ and GFP^+^ cells indicate respectively the presence of CCR2^+^ blood-borne monocyte-derived macrophages and CX3CR1^+^ macrophages. Staining with DAPI reveals nuclei, immunostaining with PECAM-1-specific antibodies reveals endothelial cells. Scale bar, 50 μm; magnified regions, scale bar 30 μm. **b)** Density of RFP^+^ (CCR2^+^) and GFP^+^ RFP^negative^ cells (CX3CR1^+^CCR2^negative^) from the choroid plexuses of the third, fourth and lateral ventricles of *CX3CR1-GFP × CCR2-RFP* mice at different disease stages. Analysis includes 3 healthy controls, 2 mice at preclinical weight loss, 7 at clinical onset, 7 at symptomatic peak, 3 at clinical remission. Data represented as mean ± standard deviation.

Secondly, we investigated whether the choroid plexus is a trafficking site for functionally committed monocyte-derived cells as suggested by our *in vitro* experiments. Thus, we visualized the *in vivo* expression of the signature pro- and anti-inflammatory enzymes iNOS and arginase-1 in the periventricular areas of *iNOS-Tomato × Arginase-EYFP* mice upon induction of EAE. In this reporter model, CCR2^+^ M^iNOS^, M^Arginase^ and M^iNOS/Arginase^ cells invade the CNS parenchyma and its interfaces in a compartmentalized manner during inflammation^25^. Confocal analysis of thick brain sections revealed a low number of arginase-1^+^, iNOS^+^arginase-1^+^ and iNOS^+^ (**Fig. 12a**) polarized macrophages at different time points of disease, starting at the weight loss preclinical phase (**Fig. 12b**). Conversely, analysis of meningeal inflammation in the surrounding ventricular walls (at clinical onset of EAE, **Fig. 12c**) confirmed a significant presence of M^iNOS^, M^Arginase^ and M^iNOS/Arginase^ cells in the adjacent inflamed brain^25^.

**Figure 12.**
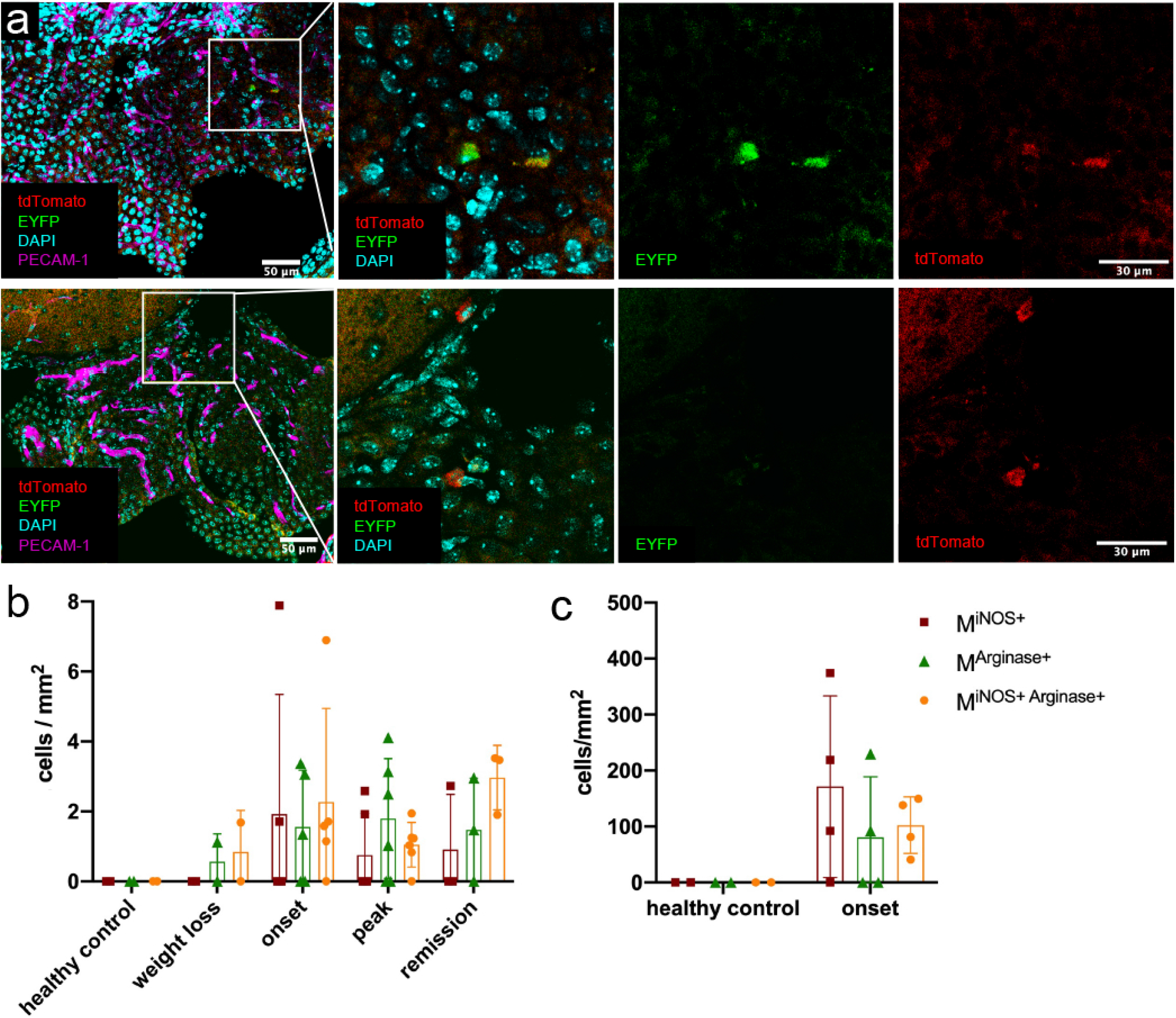
Macrophage density in the choroid plexuses of *iNOS-Tomato × Arginase-EYFP* mice during EAE. **a)** Representative confocal images of the choroid plexus from the third ventricle of a healthy *iNOS-Tomato × Arginase-EYFP* mouse (upper panel) and from the fourth ventricle of an *iNOS-Tomato × Arginase-EYFP* mouse induced with EAE (symptomatic peak, lower panel). tdTomato indicates M^iNOS^, EYFP M^Arginase^ macrophages. Staining with DAPI reveals nuclei, immunostaining with PECAM-1-specific antibodies reveals endothelial cells. Scale bar, 50μm; magnified regions, scale bar 30 μm. **b)** Density of tdTomato^+^ M^iNOS^, EYFP^+^ M^Arginase^ and tdTomato^+^EYFP^+^ M^iNOS/Arginase^ cells from the choroid plexuses of *iNOS-Tomato × Arginase-EYFP* mice at different disease stages (analysis includes 2 healthy controls, 2 mice at weight loss, 5 at clinical onset, 6 at symptomatic peak, 3 at disease remission), and **c)** from the meninges of ventricular walls at onset of EAE (analysis includes 2 healthy controls and 4 mice at clinical onset of disease). Data represented as mean ± standard deviation.

Altogether, *in vivo* analysis of brain periventricular areas reveals that functionally committed macrophages can be found in the choroid plexus during autoimmune neuroinflammation. Our data suggests that, contrary to the BBB, the BCSFB could in principle allow accumulation and CNS infiltration of pro- and anti-inflammatory macrophages during auto-aggressive CNS conditions.

## DISCUSSION

Studying the migration of immune cells through distinct CNS access gateways has considerably increased the efficacy of MS treatments^47^, but research in this field has largely focused on different classes of T and B lymphocytes^48–50^. While recent data have highlighted the prominent heterogeneity of monocyte-derived cells at CNS borders^2–4^, the trafficking routes of circulating monocytes infiltrating the CNS during neuroinflammation remain to this date surprisingly unclear.

Tissue inflammation drives the mobilization of monocytes from the bone marrow and from splenic secondary reservoir into the bloodstream^15,51^, from which these cells can potentially access the CNS at the level of the BBB, the BCSFB, or at the subarachnoid vasculature within the leptomeninges^6^. Following CNS invasion, recent investigations in the EAE neuroinflammatory model have shown that detrimental M^iNOS^ and beneficial M^Arginase^ cells differently distribute in distinct CNS anatomical compartments^25^. Can the acquisition of a pro- or anti-inflammatory expression program facilitate monocyte migration at these separate CNS gateways?

In this study, we use bone marrow-derived M^unpolarized^, M^LPS+IFNy^ and M^IL-4+IL-13^ cells to model the *in vivo* functional features of monocyte-derived cells and assess their interaction with *in vitro* BBB endothelial cells and BCSFB epithelial cells. Our overall data show that functional macrophage polarization significantly affects the adhesion and migration of macrophages across distinct CNS barriers. At the BBB, adhesion of M^LPS+IFNy^ and M^IL-4+IL-13^ polarized cells to endothelial cells was significantly reduced compared to M^Unpolarized^ cells, both in the absence and presence of physiological shear flow. Stimulation of endothelial cells with inflammatory cytokines increased both adhesion and diapedesis of M^unpolarized^ cells, indicating that neuroinflammatory conditions can increase the extravasation of monocyte-derived cells as shown for other immune cells^44,52,53^. While *in vitro* inflammatory conditions decreased pMBMEC TEER implying impaired junctional barrier properties, immune staining did not reveal any significant changes in junctional continuity, in line with the notion that BBB physical disruption is not strictly needed for cell extravasation^54^. Among the factors contributing to the reduced adhesion and migration of M^LPS+IFN-γ^ cells, the observed downregulation of *Ccr2* might play a primary role. CCL2 is a key factor in the recruitment of inflammatory monocytes and is shuttled from the abluminal to the luminal side of endothelial cells thus directly interacting with rolling immune cells^55^. This chemokine is also produced by pMBMECs^55^, altogether mediating context-dependent actions in monocyte-derived cells, including firm adhesion and arrest during trafficking^56^. Nonetheless, despite similar chemokine receptor levels and increased β2 integrin expression compared to unstimulated BMDMs, also M^IL4+IL13^ cells failed to efficiently interact with pMBMEC monolayers. While other adhesion factors and signaling molecules can obviously affect the dynamics of monocyte-derived cells at this site, the lack of antibodies specifically recognizing the morphologically activated form of integrins remain an intrinsic limitation in our approach. Overall, these data confirm our previous observations in the *iNOS-Tomato × Arginase-EYFP* model, in which analysis of cells from blood and lymph nodes did not reveal a significant amount of circulating M^iNOS^ and M^Arginase^ cells^25^. While the acquisition of a pro- or anti-inflammatory functional state is thus not a likely pre-requisite for extravasation at the BBB, subsequent macrophage accumulation in the perivascular space seems associated with a pro-inflammatory glycolytic state, as shown in both EAE and MS^25,57^. Crossing of activated endothelial cells toward the perivascular space millieu might therefore represent a key step in the priming of monocyte-derived cells toward a pro-inflammatory phenotype before invasion of the CNS parenchyma. Along this line, interaction with barrier cells was previously shown to skew monocytes into dendritic-like pro-inflammatory cells in human cultures^58^, with chemokine stimulation contributing to the complex net effect observed during neuroinflammation^59^.

Conversely, other studies have suggested that monocyte recruitment at CNS barriers other than the BBB might be associated with anti-inflammatory macrophage polarization. M^Arginase^ cells accumulate in significant numbers in the leptomeninges of *iNOS-Tomato × Arginase-EYFP* mice during EAE^25^, and anti-inflammatory macrophages were shown to traffic preferentially through the BCSFB in spinal cord injury models^60^. In our study however, while both M^unpolarized^ and M^IL4+IL13^ macrophages were able to adhere efficiently to unstimulated BCSFB epithelial cells as opposed to M^LPS+IFN-γ^ cells, all M^unpolarized^, M^LPS+IFNy^ and M^IL-4+IL-13^ cells showed comparable interaction dynamics with cytokine-stimulated pMCPECs. Moreover, despite a reduced attachment to unstimulated pMCPECs, M^LPS+IFNy^ macrophages showed an increased ability to migrate across the epithelial monolayer. Notably, both M^unpolarized^ and polarized M^LPS+IFNy^ and M^IL-4+IL-13^ cells showed increased adhesion and enhanced migration toward and through the choroid plexus epithelium compared to their interactions with pMBMECs. Taken together, these experiments suggest that the BCSFB can in principle constitute a CNS access gateway for both pro- and anti-inflammatory macrophage phenotypes.

Nonetheless, when analyzing brain sections from mice suffering from EAE, only low numbers of iNOS^+^/arginase-1^+^ macrophages could be observed *in situ*, compared to the high density of CCR2^+^ cells detected within the choroid plexus. Thus, local acquisition of an overt pro- or anti-inflammatory macrophage phenotype can happen within the choroid plexus stroma, but remains a minor phenomenon compared to the recruitment of yet-to-be-polarized monocyte-derived cells.

The BCSFB is a gateway that potentially allows blood borne cells to reach the CSF and the subarachnoid space via the foramen of Magendie, linking the fourth ventricle with the cisterna magna, or the foramina of Luschka, connecting the fourth ventricle to the cerebellopontine cisterns^61^. Once in the leptomeninges, monocyte-derived cells extensively interact with invading lymphocytes^2^, thus participating in the amplification of inflammatory processes even before reaching the parenchyma^62^. Peripheral cell migration to the CSF can however occur directly through extravasation at the level of subarachnoid vasculature^7^. For instance, transferred effector CD4^+^ T cells infiltrating the CSF during autoimmune inflammation seem to migrate through this compartment in the spinal cord^63^. Also, in an injury model, pro-inflammatory players reach the parenchyma locally through leptomeningeal vessels^60^. However, in the EAE model, the entry gateway of monocyte-derived cells accumulating in the subarachnoid space, and the mechanisms regulating a potential invasion of the CNS parenchyma crossing the pia mater and the glia limitans from this site, remain unknown.

In conclusion, our *in vitro* findings indicate that monocyte-derived cells interact more efficiently with the BCSFB compared to the BBB gateway; moreover, we could show that the BCSFB is potentially a permissive entry gateway for functionally polarized macrophages during inflammatory conditions. While our *in vitro* BMDM preparations only partially represent the complexity of *in vivo* phenotypic specifications, in which numerous stimuli are integrated^32,64^, the described paradigm allows modelling the extremities of the macrophage activation spectrum. Further experiments utilizing primary cells isolated from blood, spleen or bone marrow, and a more extensive characterization of cell trafficking at distinct sub anatomical compartments through intravital imaging, will be needed to confirm our findings.

## MATERIAL AND METHODS

### Animals

*C57BL/6J* mice were purchased from Janvier (Genest Saint Isle, France). *iNOS-tdTomato × Arginase EYFP* mice were kindly provided by Prof. Martin Kerschensteiner (LMU Munich, Germany); *CCR2-RFP × CX3CR1-GFP* mice were a kind gift of Dr. Israel F. Charo (UCSF, USA). Mice were housed in individually ventilated cages under specific pathogen-free conditions. Animal procedures were performed in accordance with the Swiss legislation on the protection of animals and were approved by the veterinary office of the Canton of Bern, Switzerland.

### Active experimental autoimmune encephalomyelitis (aEAE) induction

aEAE was induced in *C57Bl/6j* mice and in *iNOS-tdTomato × Arginase-YFP* and *CX3CR1-GFP × CCR2-RFP* mice by injection of myelin oligodendrocyte glycoprotein peptide 35-55 (MOG_35-55_ peptide, 200 μg per animal, Genscript, USA) and complete Freund’s adjuvant (Incomplete Freund’s Adjuvant, Santa Cruz Biotechnology, USA; supplemented with Mycobacterium Tubercolosis, Difco). Briefly, an emulsion of MOG_35-55_ and CFA was injected subcutaneously in mouse flanks and at the tail base at day 0; in addition, 300 ng of pertussis toxin (List Biological Laboratories, Campbell, CA, USA) was injected intraperitoneally at day 0 and day 2. Immunized mice were weighted and disease development scored daily according to a previously established system^65^. We defined four time points for analysis: weight loss (before clinical symptoms), day of clinical onset, symptomatic peak of disease, remission (7-8 days after disease onset).

### Brain isolation and vibratome sections

Mice were sacrificed and transcardially perfused with 2% paraformaldehyde (PFA, Merk Darmstadt, Germany) in Dulbecco’s phosphate-buffered saline (DPBS, Gibco, Paisley, UK); isolated brains were post-fixed in 2% PFA overnight and embedded in 2% low-melt agarose (Sigma-Aldrich, St. Louis, MO, USA) in DPBS. To cut the brain into coronal sections with a thickness of 100μm, we used a vibratome (VT1000S, Leica Biosystems, Muttenz, Switzerland) with a speed of 0.65 mm/s and a frequency of 80Hz. The samples were collected in ice cold DPBS.

### Immunofluorescence staining

Vibratome-cut free-floating slices were initially washed with 1x Tris-Buffered saline (TBS), containing 10x TBS-50mM Trizma Base (Sigma-Aldrich, St. Louis, MO, USA), 150mM NaCl (Sigma-Aldrich, Buchs, Switzerland), 1mM CaCl_2_ × 2H_2_O (Sigma-Aldrich, St. Louis, MO, USA), and ultrapure water, pH 7.4. We blocked unspecific antibody binding with blocking buffer containing TBS with 5% skimmed milk (Rapilait, Migros, Switzerland), 0.3% Triton X-100 (Sigma-Aldrich, St. Louis, MO, USA) and 0.04% NaN_3_ (Fluka Chemie, Buchs, Switzerland), pH 7.4, for 2h at room temperature (RT). To stain the blood vessels within the choroid plexus, we made use of the MEC13.3 antibody (anti-platelet endothelial cell adhesion molecule, PECAM-1/CD31, rat IgG2a, home-made), prepared in blocking buffer and incubated overnight at 4°C on a rocker. After rinsing 3 × 5 min with 1x TBS, a secondary Cy5-conjugated AffiniPure donkey anti-rat IgG (H+L) (1:200, stock of 0.5 mg/ml, Jackson ImmunoResearch Laboratories, West Grove, PA, USA, catalog number 712-175-153) antibody was applied, diluted in blocking buffer for 2h at RT. Slices were incubated with DAPI (1:5000, 1mg/ml stock, AppliChem, Darmstadt, Germany), diluted in 1x TBS for 20 min at RT. After drying, the slices were mounted with Mowiol 4-88 (water soluble hydrocolloid mucoadhesive based on polyvinyl alcohol, Sigma-Aldrich, St Louis, MO, USA).

### Cell quantification and density analysis

Z-stacks of brain slices containing choroid plexuses from the 3^rd^, 4^th^ and lateral ventricles were acquired at the LSM800 confocal microscope (Zeiss) with 40x magnification, and analyzed using Fiji (National Institute of Health, Bethesda, MD, USA). Three different z planes were chosen for analysis per slice, and area of the choroid plexuses and ventricular meninges calculated. Brain sections from *C57Bl/6j* mice (both healthy and at different time-points following EAE induction) were used to infer tissue background and exclude fluorescence artifacts. In sections from *iNOS-Tomato × Arginase-EYFP* mice, expression of tdTomato (driven by iNOS) and EYFP (driven by arginase-1) was assessed manually and blindly. In sections from *CX3CR1-GFP × CCR2-RFP* mice, numbers of RFP (driven by CCR2) and GFP (driven by CX3CR1) expressing cells was assessed automatically with a Fiji macro (**Supplementary Data 1**). The cell density in choroid plexuses from the 3^rd^, 4^th^ and lateral ventricles of each brain was calculated and extrapolated to cells per mm^2^.

### BMDM isolation and cell culture

The isolation, culture and stimulation of BMDMs were performed according to previously established protocols^25^. Briefly, mouse bones from pelvis and the hind limbs (tibia, femur) were isolated from six to twelve weeks old *C57BL/6j* male mice. The bone marrow was flushed using RPMI with glutamine (Gibco, Paisley, UK) supplemented with 10% heat inactivated fetal bovine serum (FBS, Gibco, Paisley, UK) and 100IU/ml penicillin-streptomycin (Gibco, Paisley, UK) (henceforth called BMDM media). Following filtration through 100μm filters, cells were incubated in 1ml of Ack lysing buffer (Gibco, Grand Island, NY, USA) for 5 min on ice to deplete erythrocytes. After washing, cells were cultured in BMDM media supplemented with 5ng/ml recombinant mouse macrophage colony stimulating factor (mCSF, R&D Biosystems, Minneapolis, USA, 416-ML-500) for seven days at 37°C, 5% CO_2_, at a confluence of 2 million cells/ml, in non-treated tissue culture 100mm Petri dishes (Greiner Bio-One, St. Gallen, Switzerland). At culture day seven, macrophages were polarized for 48h towards a pro-inflammatory profile (M^LPS+IFN-γ^), with 100ng/ml lipopolysaccharide from *salmonella enterica serotype typhimurium* (LPS, Sigma-Aldrich, St. Louis, MO, USA, catalog number L4516) and 10ng/ml recombinant murine IFN-γ (PeproTech, Rocky Hill, NJ, USA, catalog number 315-05); an anti-inflammatory phenotype (M^IL-4+IL-13^), with 10ng/ml recombinant murine IL-4 (R&D Biosystems, Minneapolis, USA, catalog number 404-ML) and 10ng/ml IL-13 (R&D Biosystems, Minneapolis, USA, catalog number 413-ML-025); or were left unstimulated (M^unpolarized^) in BMDM medium containing mCSF. BMDMs were collected by 10min incubation in 0.05% Trypsin (Gibco, Paisley, UK, reference 25300-054) at 37°C.

### mRNA Isolation

To isolate mRNA from BMDMs, cells were washed with ice cold sterile 1x DPBS and incubated for 5 min in 1ml TRIzol (Invitrogen, Leicestershire, UK) at RT. After cell scraping, a volume of 250μl chloroform (Merck, Darmstadt, Germany) was added. The cells were vigorously shaken for 15 seconds, left at RT for 5min, and centrifuged at a speed of 9300 × g for 15 min at 4°C. This led to the formation of a top layer aqueous phase (containing RNA), a white middle layer (containing DNA) and an organic phase at the bottom (containing proteins and organelles). The RNA phase was carefully extracted and mixed gently with 500μl isopropanol (Grogg Chemie, Stettlen, Deisswil, Switzerland). After 5min incubation and 20min 13400 × g centrifugation at 4°C, the pellet was washed once with ice cold 75% ethanol (Merck, Darmstadt, Germany). After complete ethanol evaporation, the pellet was dissolved in 25μl ultrapure water and the mRNA concentration measured using a NanoDrop (Thermo Fisher Scientific, Rochester, NY, USA). mRNA purity was assessed using the 260/280 nm and 260/230 nm ratios and the samples were used with a purity range of 1.8 to 2.

### cDNA synthesis and RTqPCR

To synthesize the DNA from single stranded mRNA via reverse transcription, we used SuperScript™ III Reverse Transcriptase cDNA synthesis kit (Invitrogen, Carlsbad, CA, USA, 18080-051) according to manufacturer’s instructions. Briefly, 700ng mRNA were incubated with 50ng random hexamers and 1mM dNTP Mix for 5min at 65°C. The reaction was then allowed to cool on ice for 5min. Afterwards, a cDNA synthesis mix containing 10x first-strand buffer, 0.1M DTT, 40IU/μl RNAseOUT (Invitrogen, Carlsbad, CA, USA, catalog number 10777-019), and 200IU/μl reverse transcriptase solution was added. The complementary DNA was catalyzed by incubating the samples at 25°C (10min) and 50°C (50min) in a PCR thermal cycler (Mastercycler X59s, Eppendorf, Hauppauge, NY, USA). The reaction was inactivated at 70°C for 15 min and allowed to cool at 4°C for 10min. To perform real time quantitative PCR reaction (RTqPCR), we used Taykon™ Low Rox SYBR MasterMix dTTP blue (Eurogentec, Liege, Belgium) according to manufacturer’s instructions. A total amount of 8.6ng cDNA per RTqPCR reaction was used. Each sample was tested in triplicates in MicroAmp™ Optical 384-well reaction plates (Applied Biosystems, Waltham, Massachusetts, USA) using the Fast Real-Time PCR System 7500 (Applied Biosystems, Waltham, Massachusetts, USA). Primer sequences are shown in **Supplementary Table 1** and the average CT values for each gene from three independent experiments are provided in **Supplementary Table 2**. We used hypoxanthine phosphoribosyltransferase (*Hprt*) as reference gene for all experiments performed. To determine the relative change in gene expression of treated samples relative to untreated control, we calculated the mean cycle threshold (Ct) value from triplicate samples of each condition for each gene, and the 2^−ΔΔCt^ value was determined by the following formula: 2^−ΔΔCt^ = 2 ^− Treated Sample * (Mean Ct value Gene of interest −Mean Ct value Reference gene) − Control Samples (Mean Ct value Gene of interest −Mean Ct value Reference gene)^.

### pMBMECs isolation and culture

pMBMECs were isolated from the cortex of six to twelve weeks old *C57BL/6j* male mice according to a previously established protocol^28^. After isolation, the cells from one brain were plated in two Matrigel-(Corning, New York, USA, reference 356231) coated 35mm dishes (ibidi GmbH, Munich, Germany) on a surface of 6.6 mm^2^ for live cell imaging experiments or in three filters of 0.5μm pore size and 6mm diameter (Corning, New York, USA, reference 3421), coated with laminin from Engelbreth-Holm-Swarm murine sarcoma basement membrane (Sigma-Aldrich, St. Louis, MO, USA) and Matrigel for transmigration experiments. The cells were grown for seven days at 37°C, 10% CO_2_ in culture media containing Dulbecco’s modified eagle medium (DMEM, Gibco, Paisley, UK) supplemented with 20% FBS (Biowest, Nuaille, France), 2% sodium pyruvate (Gibco, Paisley, UK, reference 11360-039), 2% MEM non-essential amino acids (MEM NEAA, Gibco, Paisley, UK, reference 11140-035), 50ug/ml gentamycin (Gibco, Paisley, UK, reference 15710-049) and 1ng/ml basic fibroblast growth factor (Sigma-Aldrich, St. Louis, MO, USA, F0291). For the first 48h, the media was supplemented with 4ug/ml puromycin (Sigma-Aldrich, St. Louis, MO, USA, catalog number P9620) to prevent pericyte contamination. At culture day six, cells were stimulated for 16h-20h with 20ng/ml recombinant murine IL-1β (PeproTech, Rocky Hill, NJ, USA, catalog number 211-11B-10UG) or with 5ng/ml recombinant murine TNF-α (PeproTech, Rocky Hill, NJ, USA, catalog number 211-11B-10UG) + 100IU/ml IFN-γ (PeproTech, Rocky Hill, NJ, USA, catalog number 315-05).

### pMCPECs isolation and culture

pMCPECs were isolated, cultured and stimulated according to a previously established protocol^30^ with minor adjustments. Specifically, six to twelve weeks old *C57BL/6j* male mice were sacrificed and the choroid plexuses from the lateral and fourth ventricles isolated using a stereomicroscope. Following 30 min digestion at 37°C in 1x DPBS (Gibco, Paisley, UK) containing 0.1mg/ml pronase (Roche Mannheim, Germany), epithelial cells were mechanically and enzymatically disaggregated from the choroidal structure using 0.025% trypsin-EDTA (Gibco, Paisley, UK) containing 12.5ug/ml DNAse I (Roche, Mannheim, Germany). After stopping the enzymatic reaction, the cell suspension was resuspended in pMCPECs media containing DMEM/F12 1:1 (Gibco, Paisley, UK), FBS 10% (Gibco, Paisley, UK), 2mM glutamine (Gibco, Paisley, UK), 50ug/ml gentamycine (Gibco, Paisley, UK) and plated for 2h in non-coated 35mm petri dishes (PD) (BD Biosciences, Franklin Lakes, NJ, USA) at 37°C. This step allows the detachment of fibroblast and macrophages from the choroid plexus epithelial cells. The cells were plated in pMCPECs media on 50ug/ml laminin (Roche, Mannheim, Germany) coated inverted filters of 5μm pore size and 6mm diameter (Corning, New York, USA, reference 3421) for 48h, following which the filters were placed in a 24well plate (Thermo Fisher Scientific, Rochester, NY, USA). To obtain a single monolayer, pMCPECs media supplemented with 5μg/ml human insulin (Sigma Aldrich, St Louis, MO, USA), 10ng/ml hEGF (Peprotech, Rocky Hill, NJ, USA), 2μg/ml hydrocortisone (Sigma, Buchs, Switzerland) and 20μm cytosine arabinoside (Ara-C; Sigma, St. Louis, MO, USA) was placed only below the filter, in contact with the cells. The apical side of the insert remained dry to prevent the formation of double layer culture. At culture day six, epithelial cells are stimulated whether with 10ng/ml TNF-α (PromoCell, GmbH, Heidelberg, Germany) or with 100IU/ml IFN-γ (PeproTech, Rocky Hill, NJ, USA). Unstimulated epithelial cells were used as controls.

### Live cell imaging

*In vitro* live cell imaging of BMDMs interaction with pMBMECs was performed as previously described^44^. M^unpolarized^, M^LPS+IFN-γ^ and M^IL-4+IL-13^ macrophages were resuspended in migration assay media (MAM) containing DMEM, 5% FBS, 4mM L-Glutamine (Gibco, Paisley, UK, reference A2916801) and 25mM HEPES buffer solution (Gibco, Paisley, UK, reference 15630-056) at a concentration of 1 × 10^6^ cells/ml. A total of 2 × 10^5^ cells were used per movie. Accumulation of M^unpolarized^, M^LPS+IFN-γ^ and M^IL-4+IL-13^ macrophages on pMBMECs on the flow chamber occurred for an interval of 5 min, at a low shear pressure of 0.1 dyn/cm^2^, followed by an increase in the shear flow at physiological levels of 1.5 dyn/cm^2^ for 25 min. The total recording time was 30 min, with 10 seconds interval between each frame. Image acquisition was performed using phase contrast on an inverted microscope (AxioObserver, Zeiss, Feldbach, Switzerland) with a 10x objective. The image analysis was performed using ImageJ (National Institute of Health, Bethesda, MD, USA). The number of arrested macrophages per condition was assessed at 40 seconds after onset of physiological shear flow.

### Migration assays and quantification of BMDMs

To assess macrophage migration across pMBMECs or pMCPECs monolayers under static condition, we used a transwell system as previously described^30,66^. pMBMECs and pMCPECs cultured on filters as described above were used at culture day seven after 16h cytokine stimulation, whereas BMDMs were used after 48h cytokine stimulation. Following trypsinization from culture plates, BMDMs were labelled with 1μm CellTracker™ green dye (CMFDA, Invitrogen, Rockford, IL, USA, catalog number C2925) at 37°C for 30 min. After labelling, BMDMs were washed twice with 1x DPBS and resuspended in MAM. For each pMBMEC or pMCPEC condition, 2×10^5^ BMDMs resuspended in 100μl MAM were added on the upper side of the filter. Underneath the filter, 600μl MAM were added. Laminin-coated empty filters were used as controls. BMDMs were allowed to migrate across endothelial, epithelial or empty filters for a period of 8h at 37°C, 10% CO_2_. At the end of the experiment, migrated BMDMs were collected from the bottom compartment and CMFDA^+^ cells counted using an Attune NxT flow cytometer (Thermo Fisher Scientific, Rochester, NY, USA). Later, the apical and basolateral sides were gently washed three times with 1xDPBS and the filters were 1% PFA fixed and stained with polyclonal rabbit anti-zona occludens-1 (ZO-1) antibody (Invitrogen, Rockford, IL, USA, catalog number 61-7300) to distinguish endothelial monolayer, or with monoclonal mouse anti-human E-cadherin (BD Biosciences, Franklin Lakes, NJ, USA, catalog number 610182), to delineate the epithelial layer (see below). Following fixation and staining, using a confocal microscope (LSM800 Zeiss, Germany), we acquired 20μm z-stack images starting from the upper side of each filter, with a 2μm interval. For each filter, we acquired the z-stack images at five different locations – fields of view (FOV) (upper left, upper right, center, lower left, lower right), to sample the entire filter area. After image acquisition, using ImageJ we quantified the number of macrophages attached on the upper side of the filters (corresponding to the luminal side of pMBMECs, and to the basolateral side of inverted pMCPECs), using a custom-made macro (**Supplementary figure 1**). We confirmed correct macro function by manual quantification of at least 30 fields of view from different filters. The quantification of macrophages which underwent migration across monolayers and filters (thus situated on the abluminal side of pMBMECs, or on the apical side of pMCPECs) was performed manually. For each filter, the mean and the standard deviation of the number of BMDMs from five different FOVs was used.

### Immunofluorescence staining of *in vitro* cells

pMBMECs and pMCPECs cultured on Transwell filters (Corning, New York, USA, 3421) were fixed with 1% PFA diluted in 1x DPBS for 10min at RT. After fixation and removal from the inserts, the filters were washed 3x with 1x DPBS and incubated in blocking buffer containing 5% skimmed milk (Rapilait, Migros, Switzerland), 0.3% Triton-X-100 (Sigma-Aldrich, St Louis, MO, USA), 0.04% NaN_3_ (Fluka Chemie, Buchs, Switzerland), pH 7.4 for 30min at RT. The cells were incubated afterwards in primary antibodies for 1h at RT. The following primary antibodies were used for immunofluorescence staining: polyclonal rabbit anti-ZO-1 antibody (Invitrogen, Rockford, IL, USA, catalog number 61-7300), monoclonal mouse anti-β-catenin (BD Biosciences, Franklin Lakes, NJ, USA, catalog number 610154), monoclonal mouse E-cadherin (BD Biosciences, Franklin Lakes, NJ, USA, catalog number 610182), polyclonal rabbit anti-claudin-1 (Cld-1, Invitrogen, Rockford, IL, USA, catalog number 519000), polyclonal rabbit anti-claudin-5 (Thermo Fisher Scientific, Rockford, IL, USA, catalog number 341600). Rat anti-mouse endothelial-selectin (E-Selectin, clone 10E9), rat anti-mouse intracellular adhesion molecule-1 (ICAM-1, clone 25ZC7), rat anti-mouse vascular cell adhesion molecule-1 (VCAM-1, clone 9DB3), rat anti-mouse vascular endothelial cadherin (VE-cadherin, clone 11D4) and rat anti-mouse junctional adhesion molecule-A (JAM-A, clone BV12) were used as hybridoma culture supernatants. After 3x DBPS washing steps, the samples were incubated in secondary antibodies diluted in blocking buffer for 1h at RT, under light protected conditions. The following secondary antibodies were used: alexa 488 donkey anti-mouse IgG (H+L) (Invitrogen, Eugene, OR, USA, catalog number A21202), alexa 647 goat anti-rabbit IgG (H+L) (Invitrogen, Eugene, OR, USA, catalog number A21244), Cy3-conjugated AffiniPure donkey anti-rat IgG (H+L) (Jackson ImmunoResearch, West Grove, Pa, USA, catalog number 712-165-150). Following nuclear staining with DAPI (1:5000, stock concentration of 1mg/ml, AppliChem, Darmstadt, Germany) for 5 min at RT, the filters were washed 3x with 1x DPBS, placed on glass slides (Thermo Scientific, Rochester, NY, USA) and mounted with embedding medium Mowiol (Sigma-Aldrich, St Louis, MO, USA).

### Flow cytometry

After polarization, macrophages were detached from culture plates using 0.05% trypsin/EDTA solution (Merck, Darmstadt, Germany) for 10 min at 37°C. After stopping the reaction with BMDM media, cells were washed with 1x DPBS and Fc-receptors blocked (anti-CD16/32, homemade) on ice for 15 min. The cells were then incubated with the following antibodies diluted in 1x DPBS for 30 min at 4°C, in light-protected conditions: PE-Cy7 conjugated anti-mouse CD11b (clone M1/70, BioLegend, San Diego, CA, USA, catalog number 101216); Brilliant Violet 711 conjugated anti-mouse CD45 (clone 30-F11, Biolegend, San Diego, CA, USA, catalog number 103147); fluorescein isothiocyanate (FITC) conjugated anti-mouse CD18 (β2) (clone M18/2, Invitrogen, Rockford, IL, USA, catalog number 11-0181-82); APC-efluor 780-conjugated anti-mouse CD29 (β1) (clone HMb1-1, Invitrogen, Rockford, IL, USA, catalog number 47-0291-82), alexa fluor 647-conjugated anti-mouse CD49d (α4) (BioRad, Hercules, CA, USA, catalog number MCA 1230A647T), cell viability dye eFluor 506 (Invitrogen, Rockford, IL, USA, catalog number 65-0866-14). Isotype control staining served as control. Samples were acquired using an Attune NxT cytometer (Thermo Fisher Scientific, Rochester, NY, USA). M^Unpolarized^, M^LPS+IFN-γ^ and M^IL-4+IL-13^ were gated according to their forward- and side-scattering, viability and CD11b^+^CD45^+^ expression. Analysis was performed using FlowJo™ (version 10, Ashland, OR, USA) and the relative mean fluorescence intensity (MFI) was calculated for each antibody by subtracting the MFI of antibody staining from the MFI of isotype control staining.

### Statistics

Statistical analysis was performed using GraphPad Prism 7 software (La Jolla, CA, USA). All values are presented as mean ± SD with 95% confidence interval. Asterisks indicate significant differences (*p < 0.05, **p < 0.01 and ***p < 0.001, ****p < 0.0001). One-way ANOVA with Tukey’s multiple comparison test was used for the following experiments when data was normally distributed: TEER measurements of pMBMECs and pMCPECs, BMDM integrin expression by flow cytometry, BMDM mRNA expression of inflammatory and chemokine receptors genes and quantification of CCR2^+^ and CX3CR1^+^ cells in the choroid plexus. When data did not meet the normality distribution assumption, Kruskal-Wallis Test was used instead (quantification of M^iNOS^, M^Arginase^ and M^iNOS/Arginase^ cells in the choroid plexus). Two-way ANOVA with Tukey’s multiple comparison tests was used to assess statistical significance in BMDM migration assays with pMCPECs and pMBMECs, both in presence and absence of physiological shear flow.

## AUTHOR CONTRIBUTION

G.L. and D.I. designed the experiments; D.I and S.W. performed and analyzed all experiments; G.L. and D.I. co-wrote the manuscript; G.L. supervised the study.

## ACKNOWLEDGEMENTS

We thank Prof. Britta Engelhardt for critically revising the manuscript and for continuous support and insights; Dr. Urban Deutsch, Mark Liebi, Albert Witt and animal caretakers for precious help with genotyping and mouse colony maintenance; Prof. Martin Kerschensteiner for donating the *iNOS-tdTomato × Arginase EYFP* mouse line. We thank Federico Saltarin for help in designing the Fiji macro to assess numbers of CCR2^+^ and CX3CR1^+^ macrophages in brain slices. This work is supported by a Swiss Multiple Sclerosis Society grant and by Scherbarth Foundation funding awarded to G.L.

## COMPETING INTERESTS

The authors declare no competing interests.

## SUPPLEMENTARY DATA

### Supplementary Data 1

**Fiji Macro used for the automatic quantification of BMDMs attached on the luminal side of pMBMECs or on the upper side of pMCPEC filters *in vitro***

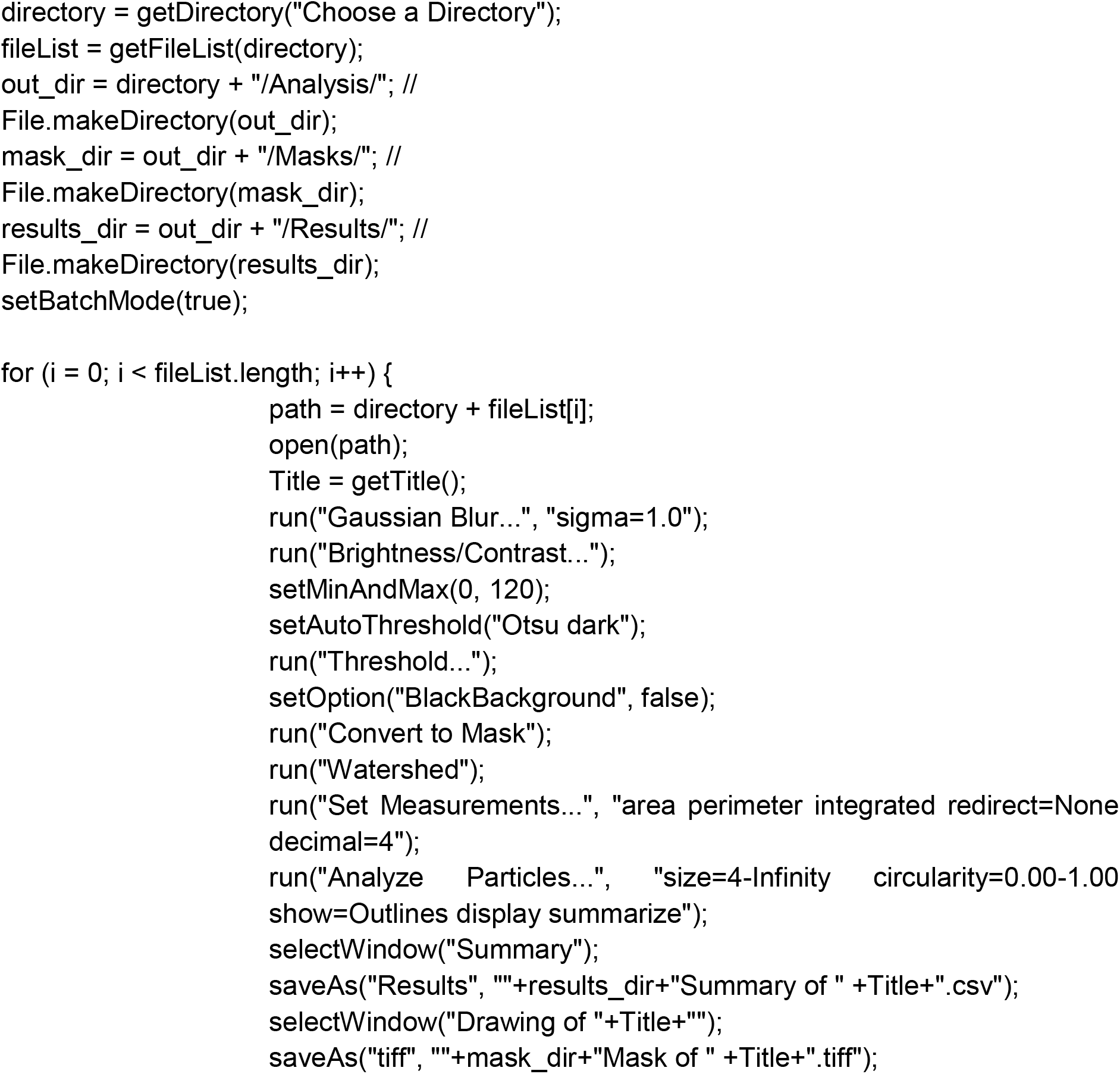

**Fiji Macro used for the automatic quantification of CX3CR1^+^ and CCR2^+^ cells in the choroid plexus of *CX3CR1-GFP × CCR2-RFP* mice**

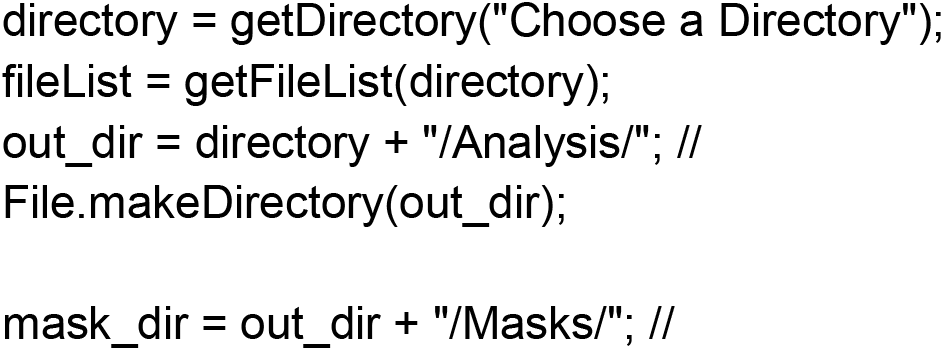
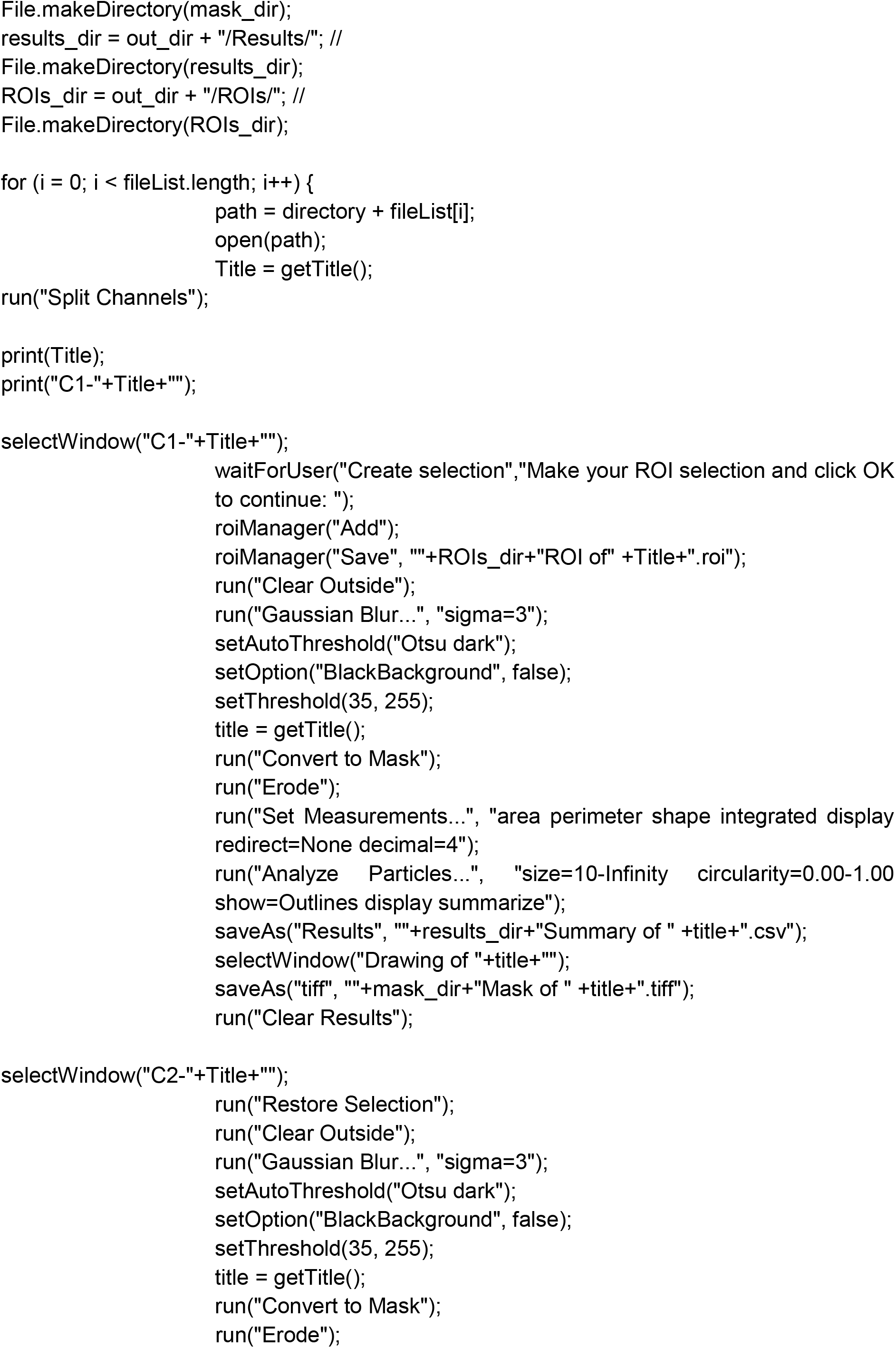
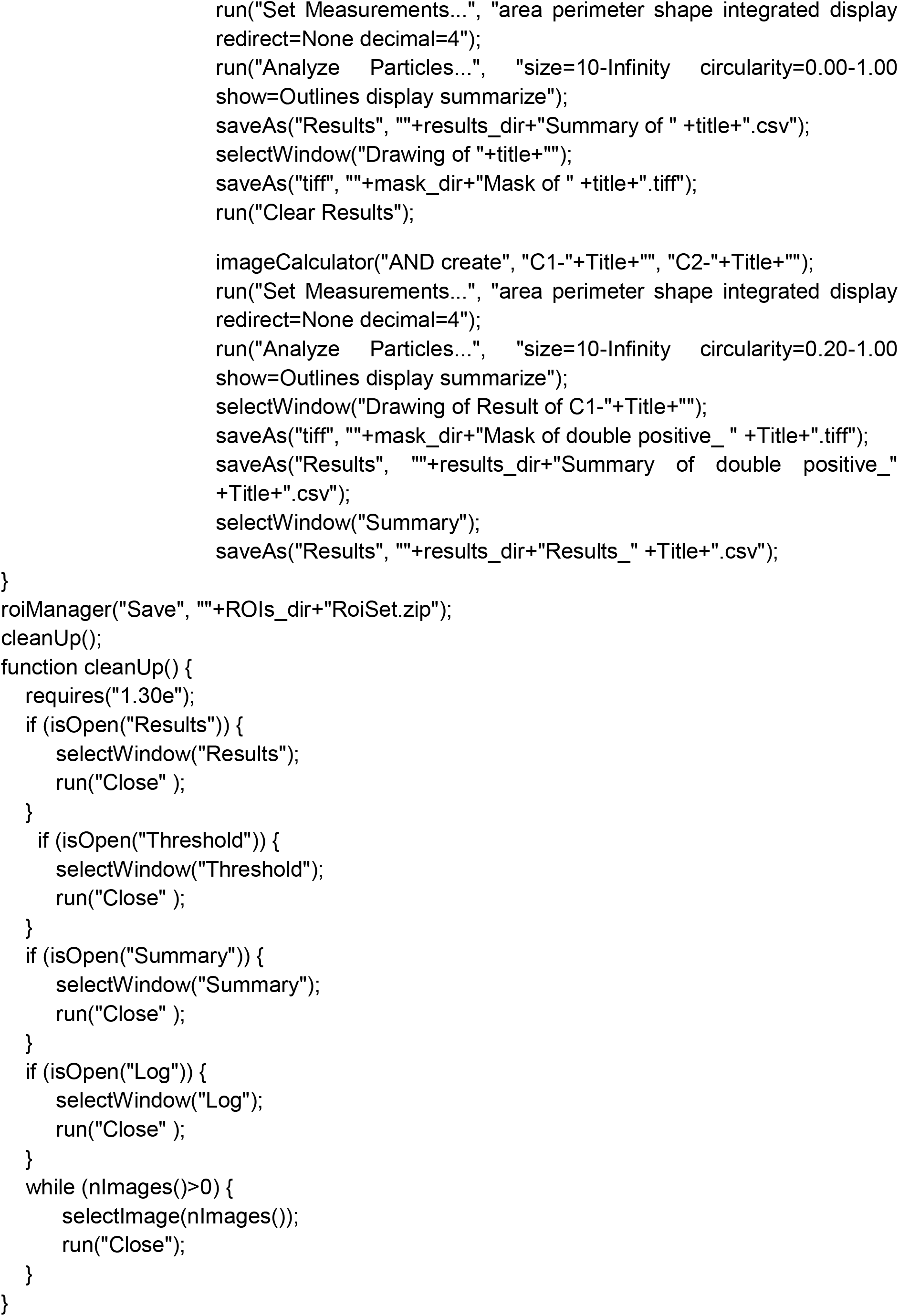

**Supplementary Table 1.**
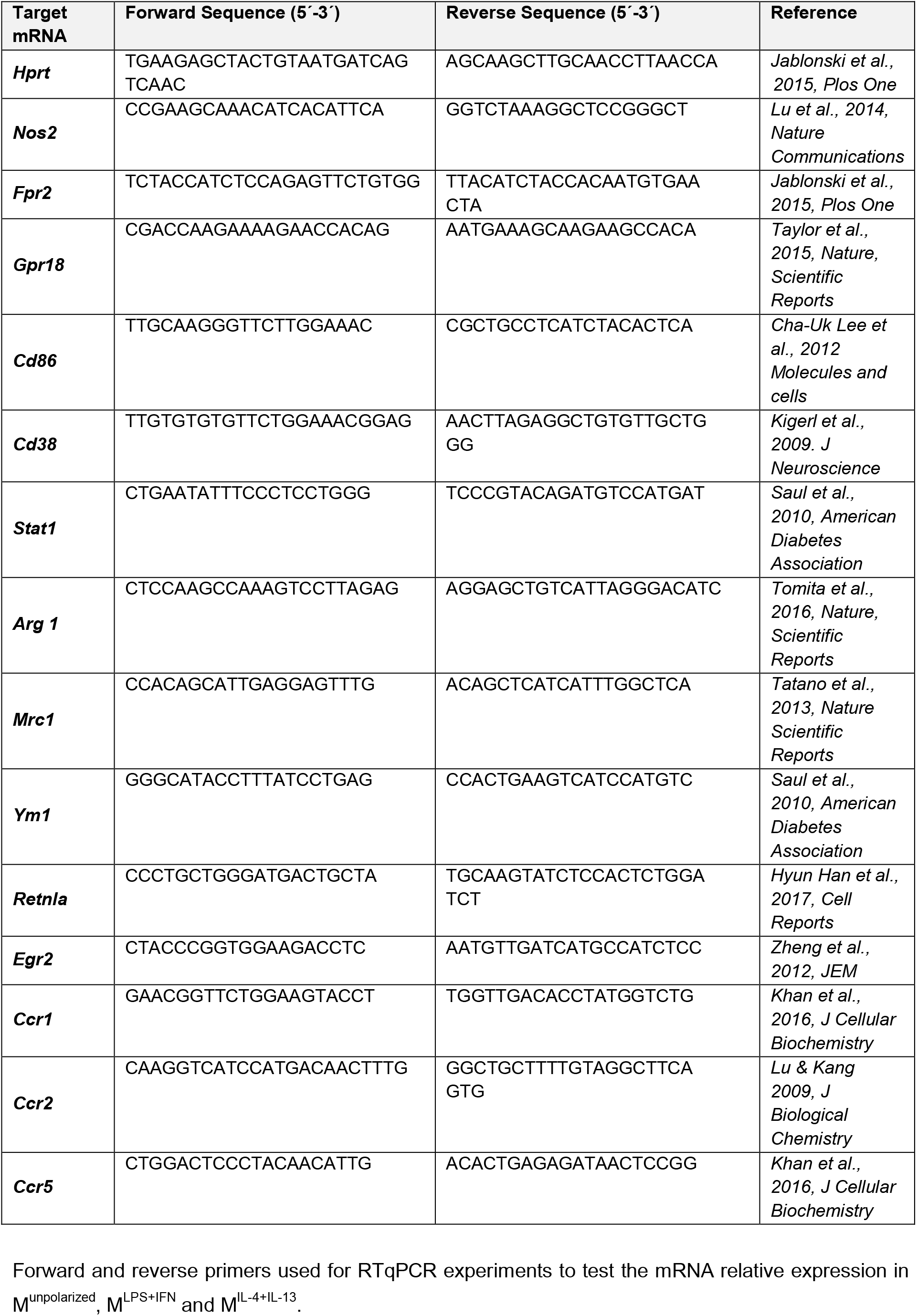

**Supplementary Table 2.**

| Target mRNA | M<sup>unpolarized</sup> |  | M<sup>LPS+IFNγ</sup> |  | M<sup>IL-4+IL-13</sup> |  | Samples Size |
| --- | --- | --- | --- | --- | --- | --- | --- |
|  | Mean Ct | STDev Ct | Mean Ct | STDev Ct | Mean Ct | STDev Ct |  |
| *Hprt* | 23.35 | 1.10 | 23.23 | 0.97 | 23.18 | 0.93 | 6.00 |
| *Nos2* | 31.42 | 3.51 | 19.10 | 2.17 | 30.55 | 1.74 | 7.00 |
| *Fpr2* | 26.99 | 0.87 | 19.46 | 0.39 | 30.31 | 0.49 | 3.00 |
| *Gpr18* | 27.93 | 1.95 | 24.59 | 0.83 | 27.49 | 0.66 | 3.00 |
| *Cd86* | 27.66 | 3.04 | 22.51 | 2.84 | 28.30 | 2.57 | 3.00 |
| *Cd38* | 30.22 | 0.61 | 22.26 | 0.80 | 30.16 | 0.45 | 3.00 |
| *Stat1* | 23.46 | 0.70 | 19.12 | 0.52 | 23.07 | 0.75 | 3.00 |
| *Arg 1* | 31.10 | 0.88 | 26.45 | 1.31 | 20.34 | 0.89 | 7.00 |
| *Mrc1* | 22.53 | 0.42 | 28.93 | 0.78 | 21.04 | 0.03 | 3.00 |
| *Ym1* | 27.65 | 0.13 | 32.24 | 0.68 | 18.85 | 1.33 | 3.00 |
| *Retnla* | 30.67 | 1.49 | 30.78 | 0.56 | 19.30 | 2.45 | 3.00 |
| *Egr2* | 23.54 | 1.48 | 27.23 | 0.21 | 21.83 | 0.92 | 3.00 |
| *Ccr1* | 24.07 | 0.38 | 23.05 | 0.46 | 23.40 | 0.54 | 3.00 |
| *Ccr2* | 23.53 | 1.32 | 28.07 | 0.99 | 24.47 | 0.84 | 3.00 |
| *Ccr5* | 20.86 | 0.25 | 20.74 | 0.05 | 20.78 | 0.28 | 3.00 |

mRNA relative expression of different pro- and anti-inflammatory and chemokine receptor genes expressed by M<sup>unpolarized</sup>, M<sup>LPS+IFN $\gamma$</sup> and M<sup>IL-4+IL-13</sup> cells following 48h stimulation. Displayed are the mean and Standard Deviation (STDev) of Cycle Threshold (Ct) values obtained from technical triplicates from six to seven experiments (for *Hprt*, *Nos2* and *Arg1*) and from three experiments (for *Fpr2*, *Gpr18*, *Cd86*, *Cd38*, *Stat1*, *Mrc1*, *Ym1*, *Retnla*, *Egr2*, *Ccr1*, *Ccr2*, *Ccr5*).

